# Anomalous Emotion Regulation & Reward Network Connectivity Underlying Suicidal & Non-Suicidal Self-Injury in Early Psychosis

**DOI:** 10.64898/2026.08.05.743099

**Authors:** Cindy An, Seema Dhaher, Melissa Kilicoglu, Jessica A. Turner, Mindy Westlund Schreiner, Aubrey Moe

**Affiliations:** Department of Psychiatry and Behavioral Health, The Ohio State University Wexner Medical Center, Columbus, Ohio, USA; Department of Psychology, The Ohio State University, Columbus, Ohio, USA; Behavioral Health, Nationwide Children’s Hospital, Columbus, Ohio, USA

**Keywords:** Early psychosis, non-suicidal self-injury, suicide, effective connectivity, fMRI

## Abstract

**Background:** Individuals with early psychosis (EP) have elevated risk for suicide, the leading cause of death in the first five years following diagnosis. Non-suicidal self-injury (NSSI) significantly predicts suicidal behavior, yet studies of self-injury often exclude participants with psychosis. We investigated effective connectivity in emotion regulation and reward network regions among participants with lifetime history of NSSI or suicide attempt (SA) with and without EP.

**Methods:** Resting-state fMRI data were acquired for 23 individuals with EP and 34 non-clinical controls (NCC). We estimated effective connectivity models for regions implicated in the self-injury literature: middle cingulate cortex (MCC), posterior cingulate cortex (PCC), caudate, putamen, posterior superior temporal gyrus (STG), orbitofrontal cortex (OFC), and insula. There were 3 models characterizing different groupings: diagnosis (NCC vs. EP); NSSI (present[+], n=21 vs. absent[-], n=36); and SA (present[+], n=21 vs. absent[-], n=36).

**Results:** EP was associated with increased STG to PCC and insula to putamen connectivity. NSSI+ (n=7 NCC, 14 EP) had increased PCC to insula lagged connectivity and increased contemporaneous bilateral putamen activity, relative to NSSI- (n=27 NCC, 9 EP). NSSI was positively correlated with lagged insula to putamen activity (p=0.016). SA and NSSI were associated with reduced PCC to caudate connectivity.

**Conclusion:** NSSI is associated with increased connectivity within emotion regulation regions and disrupted connectivity between emotion regulation and reward networks modulated by the STG and striatum. Findings are consistent with broader self-injury literature, supporting the utility of using similar interventions from other disorders to address self-injury within EP.

## 1. Introduction

Suicide is a major contributing factor to early mortality in psychotic disorders (e.g., schizophrenia, schizoaffective disorder, bipolar or depressive disorder with psychotic features), with an estimated 25-50% attempting suicide over the course of illness (Meltzer, 2001).

Individuals with schizophrenia have mortality rates 2-3 times higher and life expectancies 20 years lower than the general population (Auquier et al., 2007; Laursen et al., 2014). Suicide risk is especially elevated in the years following the initial onset of psychosis (i.e., early psychosis; EP) (S. Brown, 1997; Mortensen & Juel, 1990, 1993; Palmer et al., 2005; Robinson et al., 2010; Zaheer et al., 2020), with suicide being the leading cause of death in the first five years following initial diagnosis (Kurdyak et al., 2021). Collectively, suicidal thoughts and behaviors (STBs) and non-suicidal self-injury (NSSI) are common in psychosis, yet the neural underpinnings of these behaviors remain poorly understood.

Non-suicidal self-injury (NSSI), the deliberate destruction of one’s body tissue without suicidal intent (Nock et al., 2007), triples the likelihood of suicide deaths (Cooper et al., 2005; Lee et al., 2015). Among individuals with psychotic disorders, NSSI has a lifetime prevalence of about 33% (Lorentzen et al., 2022), and is particularly elevated during EP (Moe et al., 2022).

Although early publications of self-harm in psychosis primarily focused on case studies of extreme self-harm (Nieto et al., 1992; Waugh, 1986), recent research shows that self-harm presents similarly in psychosis relative to other psychiatric disorders (Harvey et al., 2008; Symonds et al., 2006). However, NSSI remains highly understudied in psychotic disorders.

Neural circuitry associated with STBs and NSSI include several regions such as the dorsal and ventral prefrontal cortex (PFC), cingulate, lateral temporal, orbitofrontal cortex (OFC), insula, striatum, and thalamus (Schmaal et al., 2020). Dysfunction of ventral emotional systems contributes to suicidal ideation through processes such as negative self-referential thinking, reduced positive affect, and rumination, whereas impaired dorsal cognitive control networks may contribute to suicidal behaviors through decision-making deficits. In psychosis, STBs are similarly associated with structural and functional anomalies across regions which generally overlay emotion regulation and reward networks (Aguilar et al., 2008; Besteher et al., 2016; Canal-Rivero et al., 2020, 2025; Giakoumatos et al., 2013; Girgis et al., 2021; Nanda et al., 2016; Rüsch et al., 2008; Spoletini et al., 2011; Yin et al., 2023). Previous suicide attempts (SA) among patients with psychosis are associated with blunted brain reward activity in the medial frontal gyrus (Potvin et al., 2018) and anomalies in prefrontal, temporal, and cingulate emotion regulatory activity relative to patients without SA and healthy controls (Athanassiou et al., 2021; Hoptman et al., 2024; Minzenberg et al., 2015b, 2016; Zhang et al., 2013).

Similarly, proposed models for NSSI suggest that impaired reward network reactivity contributes to trait impulsivity, leading to emotion dysregulation which increases risk for NSSI (Sauder et al., 2016). Consistent with these models, neuroimaging studies of NSSI have identified altered striatal response to rewarding stimuli (Osuch et al., 2014; Poon et al., 2019; Sauder et al., 2016), atypical frontal-limbic activation during emotion processing tasks (Demers et al., 2019; Plener et al., 2012), and altered resting-state connectivity within frontal-striatal-limbic networks (Choi et al., 2024; Cullen et al., 2020; Huang et al., 2021; Liao et al., 2024; Otto et al., 2023; Reitz et al., 2015; Santamarina-Perez et al., 2019; Vega et al., 2018; Westlund Schreiner et al., 2017; Zhou et al., 2022). However, while the neurobiology of NSSI has been studied in various transdiagnostic samples, most exclude participants with psychosis or have minimal representation despite high risk for self-injurious behaviors (SIB). Consequently, it remains unclear whether neural circuitry underlying NSSI in transdiagnostic populations generalizes to psychotic disorders. To our knowledge, the closest related study in psychosis found that in patients with schizophrenia, self-harm (unspecified whether suicidal or non-suicidal) corresponded with increased activation in the right dorsolateral PFC and left posterior cingulate cortex (PCC) during response inhibition in a go/no-go task relative to patients without self-harm (Lee et al., 2015). Overall, these studies illustrate specific brain features associated with SIB that may present across disorders and provide preliminary evidence of circuitry overlapping with psychosis neuropathology.

Prior studies of functional brain differences related to STB in psychosis have relied primarily on task-based fMRI. Though task fMRI is useful for isolating neural activity attributable to specific processes, these results can be difficult to generalize due to between-task variability. Resting-state fMRI measures intrinsic brain activity and may reflect inherent differences between populations that are less observable in task-derived data. Furthermore, prior studies focus predominantly on static connectivity, which reflects activation synchrony but not directional relationships between regions. The Group Iterative Multiple Model Estimation (GIMME) algorithm addresses this limitation by building *effective connectivity* models among pre-determined regions of interest (ROI) that characterize directional relationships over time (Gates & Molenaar, 2012). Importantly, GIMME develops network maps suiting both the individual and group, regardless of between-subject variability (Gates & Molenaar, 2012; Hillary et al., 2014). Given that psychosis and NSSI are highly heterogeneous (Keane et al., 2024; Li et al., 2026), GIMME is uniquely suited to characterize resting-state patterns in these populations.

We investigated resting-state effective connectivity differences associated with lifetime history of NSSI and SA in participants with EP and non-clinical controls (NCC). We aimed to: (1) examine effective connectivity between regions implicated in self-injury literature specific to NSSI, (2) examine effective connectivity between regions implicated in self-injury literature specific to SA, and (3) identify if psychosis-related networks overlap with circuitry associated with SIBs. We hypothesized that anomalies in effective connectivity would be associated with SIBs, and that psychosis neuropathology would contribute to increased risk for self-harm behaviors. Given limited literature, we did not make any specific hypotheses related to NSSI or SA circuitry.

## 2. Methods

### 2.1 Participants and Procedure

Data for this study were drawn from 87 participants (49 EP, 38 non-clinical controls [NCC]) from an ongoing study of neuroimaging and social cognition in EP (National Institute of Mental Health, K23MH131967, PI: Moe). The sample for the present study included only participants who completed MRI scans and passed quality control, resulting in 57 total participants (23 EP, 34 NCC). Eligibility criteria for all participants included: (1) age 18 to 30, (2) premorbid IQ estimate >70 to rule-out a primary intellectual disability, as determined by the Wide Range Achievement Test (Wilkinson & Robertson, 1993), (3) no known significant neurological abnormalities, such as seizure disorder, and (4) not meeting criteria for any active substance use disorder as determined by Structured Clinical Interview for the Diagnostic and Statistical Manual, Fifth Edition (SCID-5; First et al., 2015). SCID-5 was used to confirm primary psychotic disorder diagnoses in the EP group (i.e., schizophrenia, schizoaffective disorder, bipolar disorder with psychotic features, unspecified psychosis), and to rule out any history of psychosis in the NCC group. All EP participants were within five years of illness onset, determined as the first onset in which a participant met threshold levels of psychotic symptoms as determined by the SCID-5. All study participants provided written informed consent prior to their study participation. This study was approved by The Ohio State University Institutional Review Board.

### 2.2 Neuroimaging Data Acquisition

Neuroimaging data were collected using a Siemens Magnetom 3.0T XR Numaris/X MRI scanner. T1-weighted magnetization-prepared rapid gradient echo (MPRAGE) [TR (repetition time) = 2400.0 ms; TE (echo times) = 2.25 ms; TI (inversion time) = 1060 ms; flip angle = 8°, slice thickness = 0.8 mm; field of view (FOV) = 256 mm] images were used for registration of functional images. Eyes-open resting-state fMRI data were acquired over 10 minutes [TR = 1000.0 ms; TE = 27.00 ms; flip angle = 50°; slice thickness = 3.0 mm; FOV = 240 mm; 45 slices; voxel size = 3.0 × 3.0 × 3.0 mm], resulting in 595 whole brain volumes. A fixation cross was presented, and participants were instructed not to fall asleep. Participants were monitored to ensure that they were awake through the duration of the scan. Mock scanner usage was not standard but was available upon participant request.

### 2.3 Measures

#### 2.3.1 Suicide Risk History

All participants completed the Self-Injurious Thoughts and Behaviors Interview (SITBI) Self-Report (Nock et al., 2007) to assess for lifetime suicidal ideation, NSSI, and SA. NSSI history was defined by endorsement of any lifetime engagement in self-harm without suicidal intent (item 2). SA was defined by endorsement of any lifetime self-harm behavior in which there was suicidal intent, including interrupted and/or aborted attempts (items 3-8).

### 2.4 Analysis

#### 2.4.1 Neuroimaging Preprocessing

Neuroimaging data were preprocessed and quality checked using the Enhancing Neuroimaging Genetics through Meta-analysis (ENIGMA) Harmonized Analysis of Functional MRI pipeline (HALFpipe) v1.3.1 (Waller et al., 2022). Steps for resting-state fMRI data preprocessing followed the HALFpipe pipeline including: removal of first 5 volumes, field map correction, coregistration, spatial smoothing (full-width at half maximum (FWHM)) = 6 mm), grand mean scaling (10,000), independent component analysis-based automatic removal of motion artifacts (ICA-AROMA) based denoising, and temporal filtering (high-pass Gaussian-weighted filter width = 125.0 seconds). Slice timing correction was omitted as is recommended for multi-band fMRI data.

Quality control was conducted within HALFpipe by visually reviewing the T1w skull stripping and segmentation, T1w spatial normalization, EPI signal-to-noise ratio, ICA-based artifact removal, and spatial normalization. Participants who had mean framewise displacement (FD) > 0.5 mm were excluded from the analysis. Four participants with EP were excluded: three for high mean FD and one for missing BOLD signal, resulting in 57 subjects overall.

HALFpipe was used to segment the resting-state fMRI bold signal into regions from the Schaefer combined atlas, which includes 400 parcellations of the cerebral cortex based on Yeo (2011) 17-network atlas (Schaefer et al., 2018), 17 subcortical regions from the FreeSurfer atlas (Fischl et al., 2002), and the Buckner 17-network cerebellar atlas (Buckner et al., 2011).

#### 2.4.2 ROI Selection

We selected regions of interest by identifying commonly significant regions in neuroimaging literature on adolescent NSSI and on STBs in psychosis. (See **Supplemental Table 1a** for a full presentation). Commonly significant regions were the cingulate, insula, striatum, STG, and OFC. Despite the striatum overall being notable, the nucleus accumbens was not included in our chosen regions because most participants were missing BOLD signal due to signal dropout from that region in our dataset. The reported MNI coordinates for the selected ROIs from literature were overlayed onto the Schaefer combined atlas to identify the most suitable parcellation or aggregation of parcellations to represent the region (**Supplemental Table 2**).

For the following analyses, the regions of interest were both the right and left middle cingulate cortex (MCC), PCC, caudate, putamen, posterior STG, OFC, and middle insula, which comprise components of the emotion regulation and reward networks (**Figure 1**). For regions represented by multiple parcellations, the time series across parcellations were averaged, resulting in one time series of BOLD signal activity per ROI.

**Figure 1.**
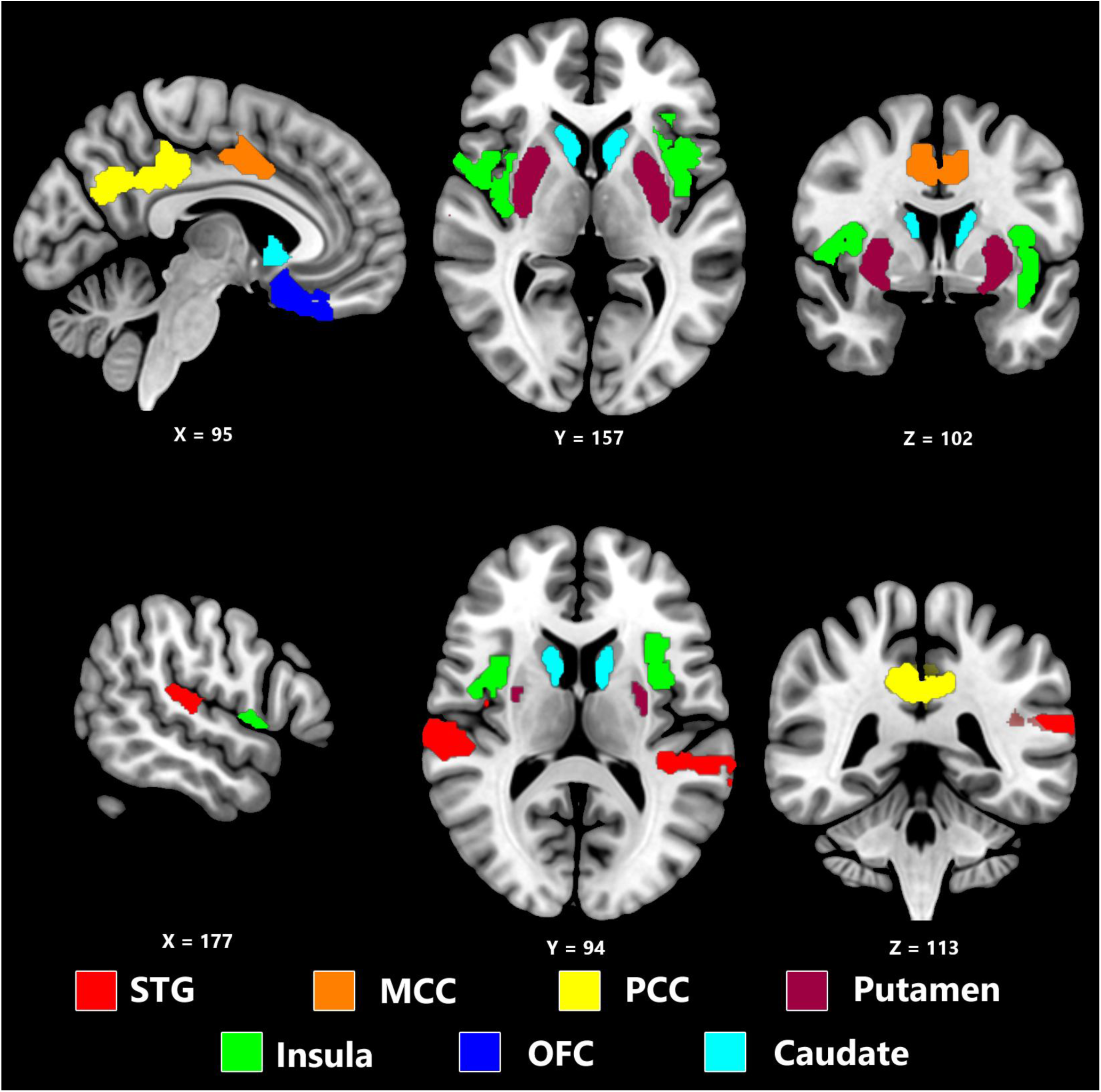
Region of interest masks. All coordinates are MNI coordinates: top (X=95, Y=157, Z=102) and bottom (X=177, Y=94, Z=113). STG = superior temporal gyrus; MCC = middle cingulate gyrus; PCC = posterior cingulate gyrus; OFC = orbitofrontal cortex.

#### 2.4.3 GIMME Modeling

Group Iterative Multiple Model Estimation (GIMME) (version 0.8.3) is a method building on unified structural equation modelling to estimate directional functional networks, or effective connectivity (Gates & Molenaar, 2012). These functional networks include autoregressive (self-predictive activity across time), contemporaneous (between region connectivity within time), and lagged (between region connectivity across a time lag of 1 TR) paths. GIMME is a unique method of measuring effective connectivity because, unlike other methods, it produces connectivity maps which capture group-level similarities despite individual-level heterogeneity within the sample. Specifically, it develops group-level models of effective connectivity which fit a majority of individuals (set to 75% for our analyses).

Supervised GIMME was used to create effective connectivity maps characterizing directed activity in the selected ROIs for each predetermined subgroup. In R (version 4.4.1), each participant’s BOLD signal time series was concatenated. A vector defined subgroup identity for each model based on clinical group, NSSI history, and SA history. The time series data frame and the subgrouping vector were input into the GIMME function from the *gimme* package (Lane et al., 2025). Standardize was set to TRUE to transform the input unstandardized time series to a mean of zero and standard deviation of one. Three models represented separate subgroupings: 1) NCC vs. EP, 2) NSSI- vs. NSSI+ (absent[-] vs. present[+]), 3) SA- vs. SA+ (absent[-] vs. present[+]). First, GIMME estimates individual-level models, starting with autoregressive paths, for each participant. This individual-level model generates autoregressive, contemporaneous, and lagged paths until excellent model fit is obtained, which is defined as meeting threshold for two of four canonical fit statistics traditionally used in structural equation modelling (Cangur & Ercan, 2015). It then estimates group-level paths (paths shared by 75% of all participants in the model). Finally, it estimates subgroup-level paths (paths shared by 75% of participants in the defined subgroup of the model).

#### 2.4.4 Path Coefficient Regressions

GIMME calculates path coefficients for all autoregressive, lagged, and contemporaneous paths, representing the strength of effective connectivity. In R, multiple regressions were conducted to assess the effect of diagnostic group and SIB (NSSI and SA in separate models) on strength of effective connectivity (standardized β) for group-level paths in each model. All models included diagnostic group, NSSI presence, and number of SA as variables of interest, and age, sex, and mean FD were included as covariates. The Benjamini-Hochberg procedure was used to correct for multiple comparisons (Benjamini & Hochberg, 1995).

#### 2.4.5 Sensitivity Analysis

Due to the small sample size, we conducted a leave-one-out inspired sensitivity analysis to assess the stability of GIMME maps. For all models, we removed one participant and generated a connectivity map following the same procedure as above. This resulted in 57 connectivity maps per model, representing connectivity patterns following one removed participant. Comparing these leave-one-out models to the original model, we determined which paths were most stable. For example, if a path from the original model remains in 50 of the 57 leave-one-out models, we could say it is ∼87% stable. These stability measures are included with our results.

### 2.5 Secondary Analysis

#### 2.5.1 Static Connectivity

Static connectivity, a more commonly used method to assess resting-state functional connectivity, represents between-ROI time series correlations. Static connectivity was also assessed using the same preselected ROIs as an additional comparison to the primary effective connectivity analysis. In R, the BOLD signal activity time series was correlated across ROIs using the function *cor* to calculate Pearson correlations, or static connectivity measures for each participant (R Core Team, 2024). These static connectivity measures were transformed from a range of [-1, 1] to [-∞,∞] using the Fisher Z-Transformation. Multiple regressions assessed the effect of subgroup identity on static connectivity strength. All regressions included age, sex, and mean FD as covariates. The Benjamini-Hochberg procedure was used to correct for multiple comparisons (Benjamini & Hochberg, 1995).

## 3. Results

### 3.1 Assessments and Subgroup Demographics

**Tables 1**, **2**, and **3** are provided to describe the demographic characteristics for each of the model subgroups. Large portions of the NSSI+ subgroup (66.7%) and the SA+ subgroup (76.2%) were composed of participants with EP.

**Table 1.**
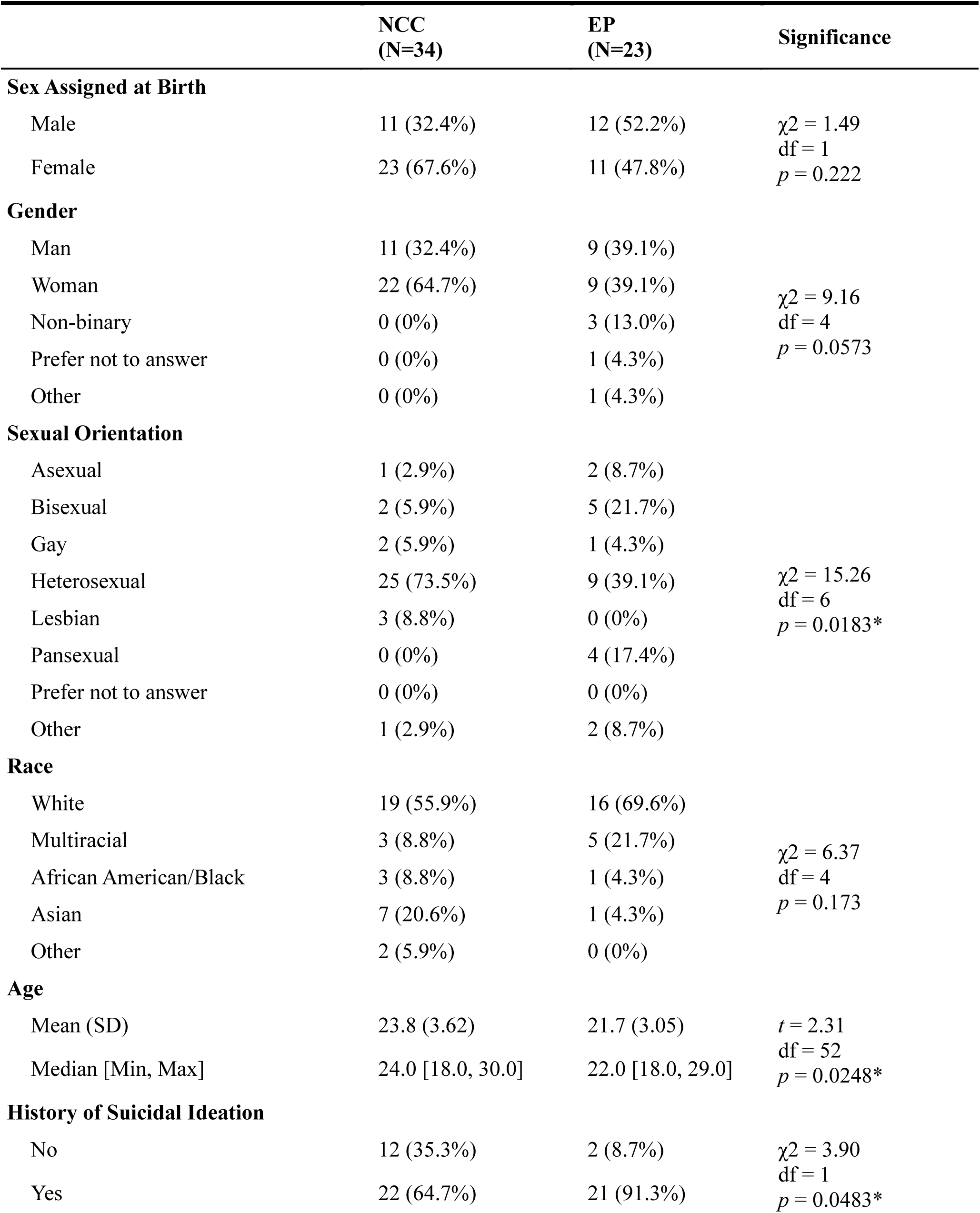
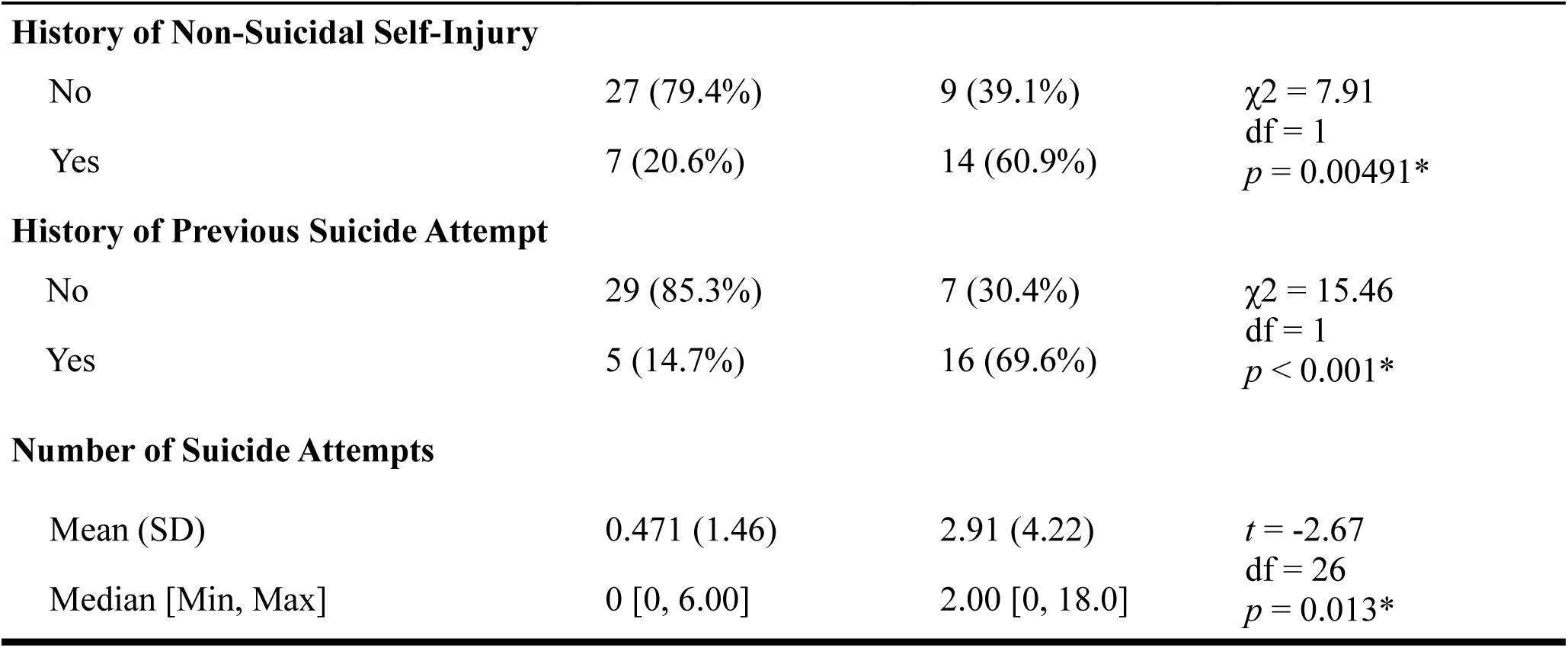
Demographic and Suicidal Thoughts and Behaviors Characteristics. Reports on suicidal ideation, non-suicidal self-injury, and suicide attempt presence are lifetime prevalence. NCC = non-clinical control; EP = early psychosis; df = degrees of freedom; * = significant at *p* < 0.05.

|  | NCC<br>(N=34) | EP<br>(N=23) | Significance |
| --- | --- | --- | --- |
| Sex Assigned at Birth |  |  |  |
| Male | 11 (32.4%) | 12 (52.2%) | $\chi^2 = 1.49$<br>df = 1<br>$p = 0.222$ |
| Female | 23 (67.6%) | 11 (47.8%) |  |
| Gender |  |  |  |
| Man | 11 (32.4%) | 9 (39.1%) | $\chi^2 = 9.16$<br>df = 4<br>$p = 0.0573$ |
| Woman | 22 (64.7%) | 9 (39.1%) |  |
| Non-binary | 0 (0%) | 3 (13.0%) |  |
| Prefer not to answer | 0 (0%) | 1 (4.3%) |  |
| Other | 0 (0%) | 1 (4.3%) |  |
| Sexual Orientation |  |  |  |
| Asexual | 1 (2.9%) | 2 (8.7%) | $\chi^2 = 15.26$<br>df = 6<br>$p = 0.0183^*$ |
| Bisexual | 2 (5.9%) | 5 (21.7%) |  |
| Gay | 2 (5.9%) | 1 (4.3%) |  |
| Heterosexual | 25 (73.5%) | 9 (39.1%) |  |
| Lesbian | 3 (8.8%) | 0 (0%) |  |
| Pansexual | 0 (0%) | 4 (17.4%) |  |
| Prefer not to answer | 0 (0%) | 0 (0%) |  |
| Other | 1 (2.9%) | 2 (8.7%) |  |
| Race |  |  |  |
| White | 19 (55.9%) | 16 (69.6%) | $\chi^2 = 6.37$<br>df = 4<br>$p = 0.173$ |
| Multiracial | 3 (8.8%) | 5 (21.7%) |  |
| African American/Black | 3 (8.8%) | 1 (4.3%) |  |
| Asian | 7 (20.6%) | 1 (4.3%) |  |
| Other | 2 (5.9%) | 0 (0%) |  |
| Age |  |  |  |
| Mean (SD) | 23.8 (3.62) | 21.7 (3.05) | $t = 2.31$<br>df = 52<br>$p = 0.0248^*$ |
| Median [Min, Max] | 24.0 [18.0, 30.0] | 22.0 [18.0, 29.0] |  |
| History of Suicidal Ideation |  |  |  |
| No | 12 (35.3%) | 2 (8.7%) | $\chi^2 = 3.90$<br>df = 1<br>$p = 0.0483^*$ |
| Yes | 22 (64.7%) | 21 (91.3%) |  |
| History of Non-Suicidal Self-Injury |  |  |  |
| No | 27 (79.4%) | 9 (39.1%) | $\chi^2 = 7.91$<br>df = 1<br>$p = 0.00491^*$ |
| Yes | 7 (20.6%) | 14 (60.9%) |  |
| History of Previous Suicide Attempt |  |  |  |
| No | 29 (85.3%) | 7 (30.4%) | $\chi^2 = 15.46$<br>df = 1<br>$p < 0.001^*$ |
| Yes | 5 (14.7%) | 16 (69.6%) |  |
| Number of Suicide Attempts |  |  |  |
| Mean (SD) | 0.471 (1.46) | 2.91 (4.22) | $t = -2.67$<br>df = 26<br>$p = 0.013^*$ |
| Median [Min, Max] | 0 [0, 6.00] | 2.00 [0, 18.0] |  |

**Table 2.** Demographic and assessment characteristics of participants without lifetime presence of non-suicidal self-injury [NSSI-, n=36] compared to participants with lifetime presence of non-suicidal self-injury [NSSI+, n=21]. df = degrees of freedom; * = significant at *p* < 0.05.

|  | NSSI-<br>(N=36) | NSSI+<br>(N=21) | Significance |
| --- | --- | --- | --- |
| **Diagnostic Subgroup** |  |  |  |
| Control | 27 (75.0%) | 7 (33.3%) | $\chi^2 = 7.91$<br>df = 1 |
| First Episode Psychosis | 9 (25.0%) | 14 (66.7%) | $p = 0.00491^*$ |
| **Sex Assigned at Birth** |  |  |  |
| Male | 17 (47.2%) | 6 (28.6%) | $\chi^2 = 1.22$<br>df = 1 |
| Female | 19 (52.8%) | 15 (71.4%) | $p = 0.269$ |
| **Age** |  |  |  |
| Mean (SD) | 23.5 (3.52) | 22.0 (3.41) | $t = 1.61$<br>df = 43 |
| Median [Min, Max] | 23.5 [18.0, 30.0] | 21.0 [18.0, 30.0] | $p = 0.114$ |
| **History of Previous Suicide Attempt** |  |  |  |
| No | 31 (86.1%) | 5 (23.8%) | $\chi^2 = 19.53$<br>df = 1 |
| Yes | 5 (13.9%) | 16 (76.2%) | $p < 0.001^*$ |
| **Number of Suicide Attempts** |  |  |  |
| Mean (SD) | 0.222 (0.591) | 3.57 (4.38) | $t = -3.49$<br>df = 20 |
| Median [Min, Max] | 0 [0, 2.00] | 3.00 [0, 18.0] | $p = 0.00226^*$ |

**Table 3.** Demographic and assessment characteristics of participants without lifetime history of suicide attempt(s) [SA-, n=36] compared to participants with lifetime history of suicide attempt(s) [SA+, n=21]. df = degrees of freedom; * = significant at *p* < 0.05.

|  | SA-<br>(N=36) | SA+<br>(N=21) | Significance |
| --- | --- | --- | --- |
| **Diagnostic Subgroup** |  |  |  |
| Control | 29 (80.6%) | 5 (23.8%) | $\chi^2 = 15.47$<br>df = 1 |
| First Episode Psychosis | 7 (19.4%) | 16 (76.2%) | $p < 0.001^*$ |
| **Sex Assigned at Birth** |  |  |  |
| Male | 16 (44.4%) | 7 (33.3%) | $\chi^2 = 0.30$<br>df = 1 |
| Female | 20 (55.6%) | 14 (66.7%) | $p = 0.586$ |
| **Age** |  |  |  |
| Mean (SD) | 23.7 (3.47) | 21.8 (3.36) | $t = 2.04$<br>df = 43 |
| Median [Min, Max] | 24.0 [18.0, 30.0] | 21.0 [18.0, 29.0] | $p = 0.0476^*$ |
| **History of Non-Suicidal Self-Injury** |  |  |  |
| No | 31 (86.1%) | 5 (23.8%) | $\chi^2 = 19.53$<br>df = 1 |
| Yes | 5 (13.9%) | 16 (76.2%) | $p < 0.001^*$ |
| **Number of Suicide Attempts** |  |  |  |
| Mean (SD) | 0 (0) | 3.95 (4.10) |  |
| Median [Min, Max] | 0 [0, 0] | 3.00 [1.00, 18.0] |  |

### 3.2 GIMME Modeling

Effective connectivity maps for the NCC vs. EP model, NSSI- vs. NSSI+ model, and SA-vs. SA+ model are respectively shown in **Figures 2**, **3**, and **4**. Group-level paths, or paths representing 75% of the entire sample, are visualized as black lines while subgroup-level paths, or paths representing 75% of the participants in the subgroup, are visualized as green lines. Solid lines represent contemporaneous paths, or correlations between regions within time, and dashed lines represent lagged paths, or correlations across a time lag of 1 TR. Regions are represented as nodes and are colored to represent the reward and emotion regulation networks. Stability measures of all paths are presented in **Supplemental Table 3**.

**Figure 2.**
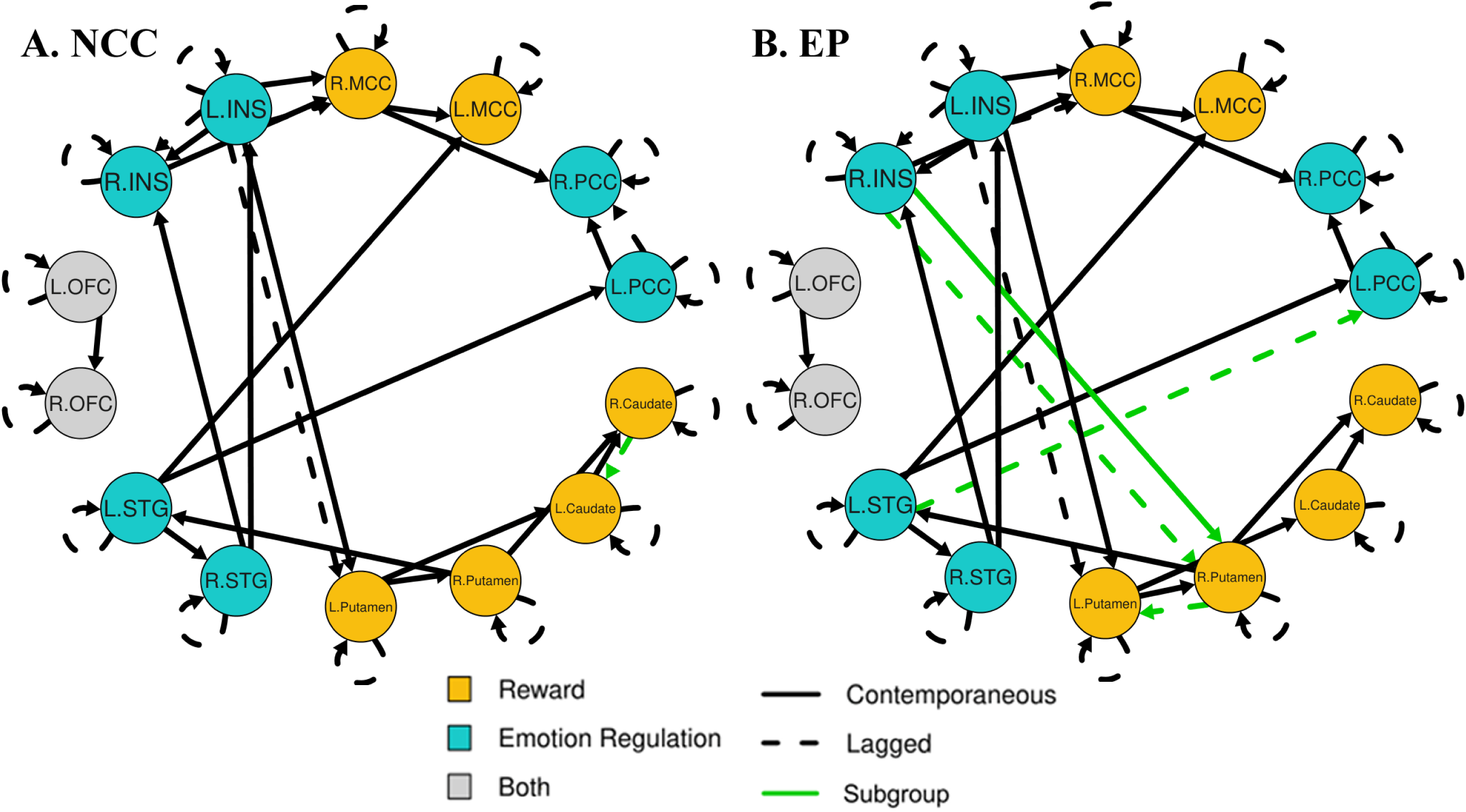
Effective connectivity maps for non-clinical controls [NCC] (A, n=34) and participants with early psychosis [EP] (B, n=23). Black lines indicate group-level paths representing 75% of participants across subgroups. Green lines indicate subgroup-level paths representing 75% of participants within a subgroup that are not present in other subgroups. L = left; R = right; MCC = middle cingulate cortex; PCC = posterior cingulate cortex; STG = superior temporal gyrus; OFC = orbitofrontal cortex; INS = insula.

#### 3.2.1 NCC vs. EP Model

In the EP effective connectivity map (**Figure 2, B**), the following subgroup-level paths emerged as uniquely characterizing EP: R.Insula → R.Putamen (85.96% stable), R.Insula → R.Putamen lagged (85.96% stable), R.Putamen → L.Putamen lagged (89.47% stable), and L.STG → L.PCC lagged (100% stable). Subgroup-level paths which uniquely characterized NCC (**Figure 2, A**) included: R.Caudate → L.Caudate lagged (98.24% stable). As a majority of the NSSI+ and SA+ subgroups were composed of participants with EP, overlapping subgroup-level paths between these models are indistinguishable from EP pathology, which itself may contribute to the development of SIBs but are not specific to self-injury neurobiology. Such paths include the R.Insula → R.Putamen contemporaneous and lagged EP paths which overlap with both NSSI+ and SA+ models and the R.Putamen → L.Putamen lagged EP path which overlaps with the SA+ model. The L.STG → L.PCC lagged EP path and the R.Caudate → L.Caudate lagged NCC path were specific to the NCC vs. EP model, inferring that increased connectivity in these lagged paths may contribute to EP circuitry separate from SIB circuitry.

#### 3.2.2 NSSI- vs. NSSI+ Model

In the NSSI+ effective connectivity map (**Figure 3, B**), the following subgroup-level paths emerged as characterizing NSSI+: R.Insula → R.Putamen (87.71% stable), R.Putamen → L.Putamen (66.66% stable), R.Insula → R.Putamen lagged (85.96% stable), and L.PCC → L.Insula lagged (73.68% stable). Subgroup-level paths which characterized NSSI- (**Figure 3, A**) included: R.PCC → L.Caudate (96.49% stable). The L.PCC → L.Insula lagged and R.Putamen → L.Putamen paths were specific to NSSI+ and did not appear in the EP or SA+ models.

**Figure 3.**
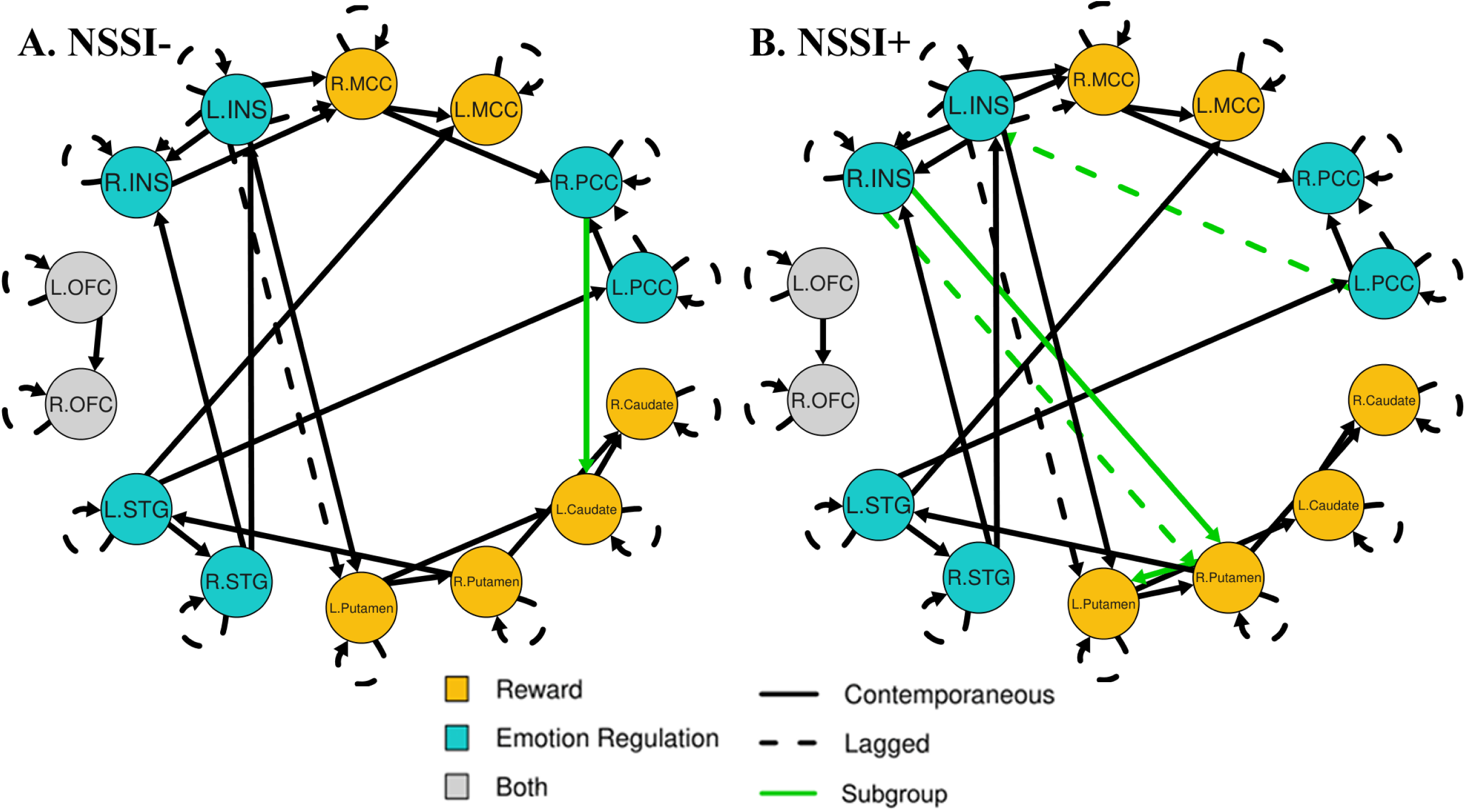
Effective connectivity maps for participants without lifetime presence of non-suicidal self-injury [NSSI-] (A, n=36) and participants with lifetime presence of non-suicidal self-injury [NSSI+] (B, n=21). Black lines indicate group-level paths representing 75% of participants across subgroups. Green lines indicate subgroup-level paths representing 75% of participants within a subgroup that are not present in other subgroups. L = left; R = right; MCC = middle cingulate cortex; PCC = posterior cingulate cortex; STG = superior temporal gyrus; OFC = orbitofrontal cortex; INS = insula.

#### 3.2.3 SA- vs. SA+ Model

In the SA+ effective connectivity map (**Figure 4, B**), the following subgroup-level paths emerged as characterizing SA+: R.Insula → R.Putamen (87.71% stable), R.Insula → R.Putamen lagged (85.96% stable), and R.Putamen → L.Putamen lagged (100% stable). Subgroup-level paths which characterized SA- (**Figure 4, A**) included: R.PCC → L.Caudate (89.47% stable). No paths were distinct to the SA model but elevated R.PCC → L.Caudate connectivity emerged continuously characterizing both SA- and NSSI-.

**Figure 4.**
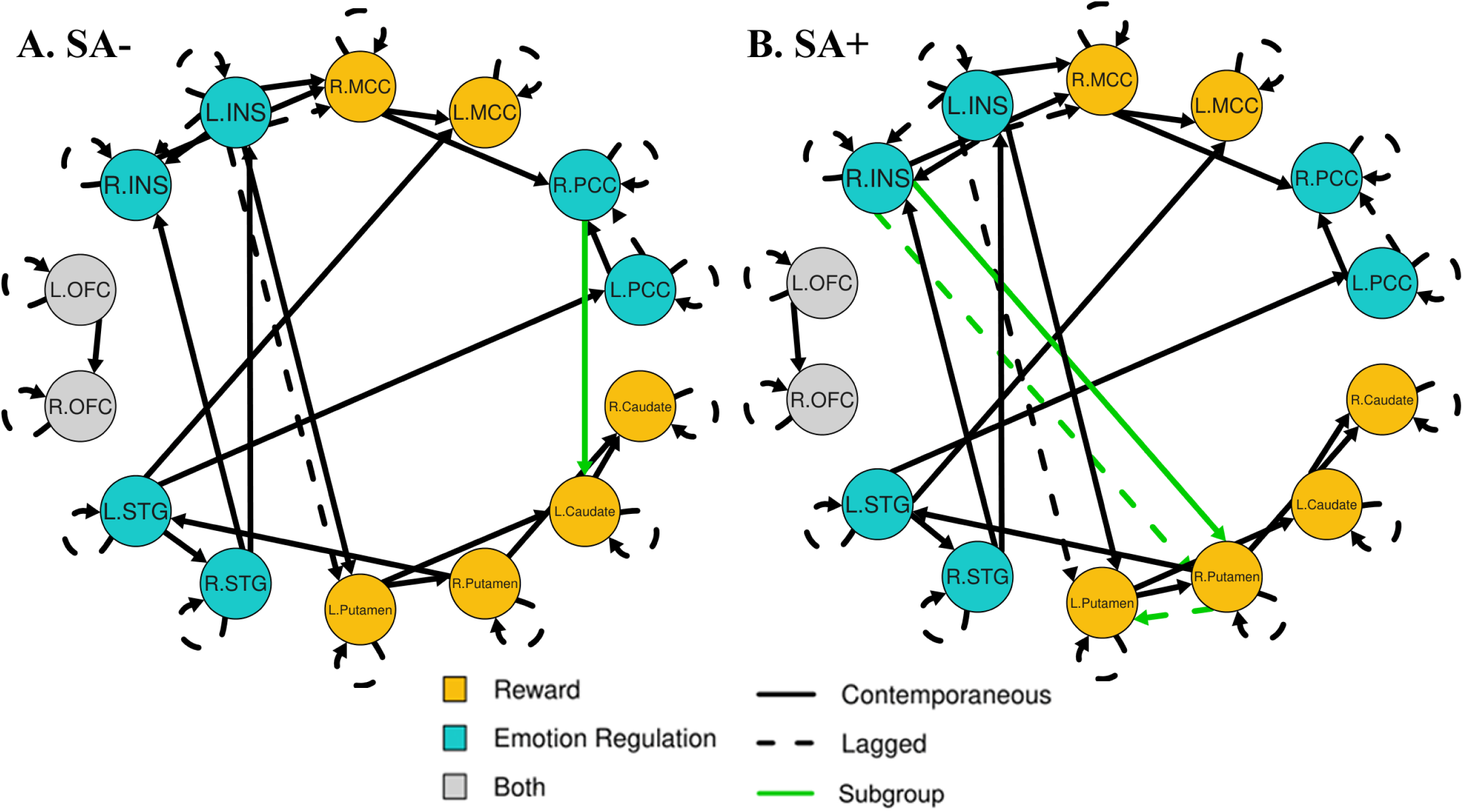
Effective connectivity maps for participants without lifetime history of suicide attempt(s) [SA-] (A, n=36) and participants with lifetime history of suicide attempt(s) [SA+] (B, n=21). Black lines indicate group-level paths representing 75% of participants across subgroups. Green lines indicate subgroup-level paths representing 75% of participants within a subgroup that are not present in other subgroups. L = left; R = right; MCC = middle cingulate cortex; PCC = posterior cingulate cortex; STG = superior temporal gyrus; OFC = orbitofrontal cortex; INS = insula.

### 3.3 Path Coefficients

A table of all significant path coefficients are provided in **Supplemental Table 4.** Path coefficients measure the strength of effective connectivity for all group-level paths. Only results which survived Bonferroni corrected *p* < 0.05 are reported. If paths were significant in multiple models, the least significant corrected *p*-value was reported, but all *p*-values are in **Supplemental Table 4**. No paths were significantly related to SA.

EP was significantly correlated with increased L.Caudate autoregressive (*p* = 0.029) and L.STG → L.PCC (*p* = 0.003) effective connectivity strength. Viewed in tandem with the subgroup-level paths from the NCC vs. EP effective connectivity map, EP exhibits overall increased striatal effective connectivity compared to NCC. This elevated striatal connectivity may drive increased STG to PCC connectivity, reflected by both elevated lagged and contemporaneous connectivity between L.STG → L.PCC. Importantly, L.STG → L.PCC effective connectivity strength was also significantly associated with FD (*p* = 0.022), limiting the confidence of interpretations.

NSSI was associated with increased strength in L.Insula → L.Putamen lagged (*p* = 0.016) and right putamen autoregressive (*p* = 0.001) effective connectivity. NSSI was also associated with reduced strength in L.Insula → L.Putamen contemporaneous (*p* = 0.009), left putamen autoregressive (*p* = 0.006), and L.Putamen → R.Putamen (*p* = 2.7E-04) effective connectivity. These results viewed alongside the GIMME models suggest that insula connectivity drives altered autoregressive connectivity in the left putamen which drives altered autoregressive connectivity in the right putamen. In NSSI, connectivity between emotion regulation and reward networks may favor lagged paths, indicating disrupted processing between networks.

### 3.4 Secondary Analyses

#### 3.4.1 Static Connectivity

Results from the regressions assessing the effect of subgroup identity on static connectivity strength are provided in **Table 4**. Static connectivity analyses were all insignificant after adjusting for multiple comparisons. Nominally significant results were found for the NCC vs. EP and SA- vs. SA+ models but did not overlap with results identified by GIMME. EP was associated with reduced bilateral putamen to bilateral MCC connectivity (uncorrected *p* = 0.040). SA+ was associated with reduced PCC and MCC connectivity (uncorrected *p* = 0.023) and OFC and caudate connectivity (uncorrected *p* = 0.049).

**Table 4.**
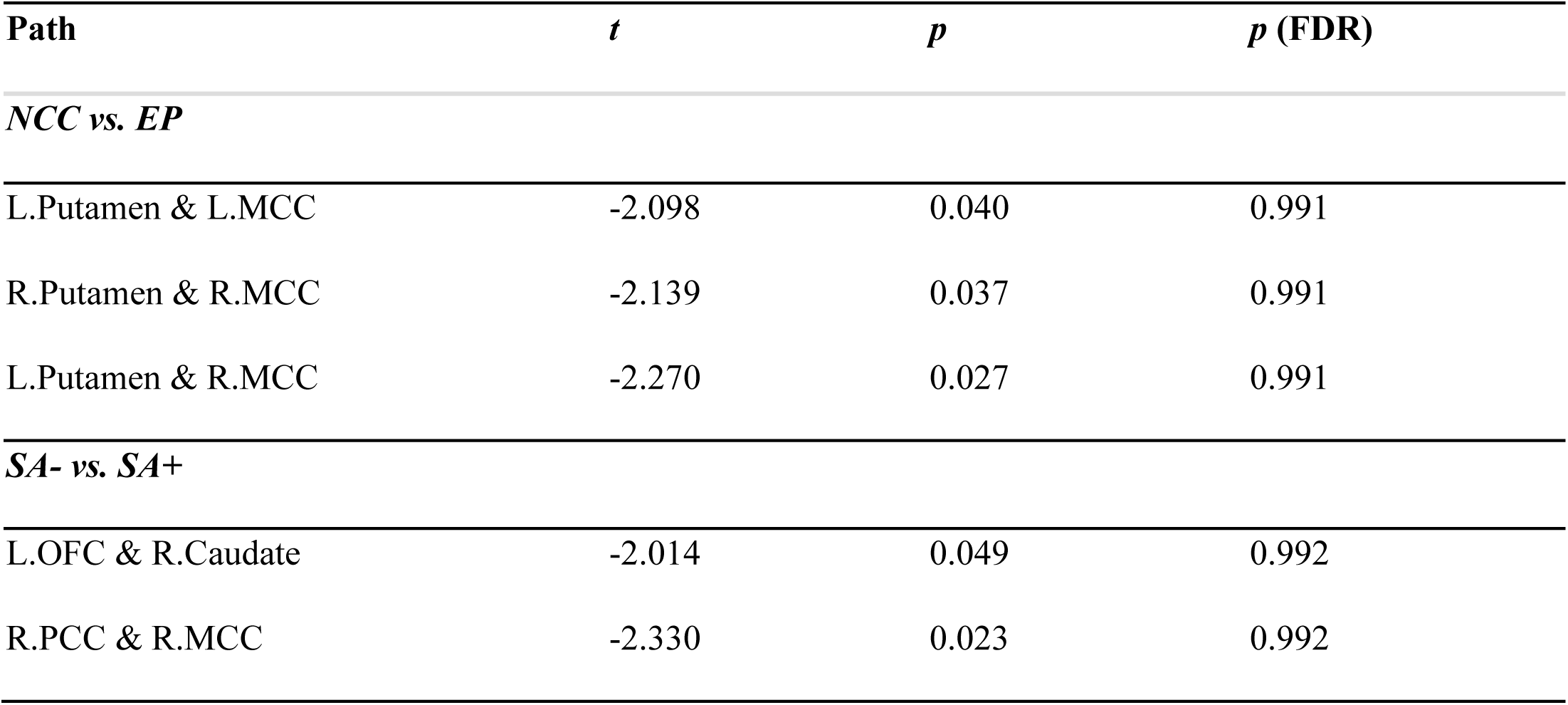
Static connectivity t-values associated with subgroup identity with significance uncorrected p < 0.05. Regressions on Fisher Z-Transformed static connectivity values included age, sex, and mean FD as covariates. Subgroup identities were coded as 1 (NCC, NSSI-, SA-) and 2 (EP, NSSI+, SA+). NCC = non-clinical control; EP = early psychosis; NSSI = non-suicidal self-injury; SA = suicide attempt; FDR = false discovery rate corrected. L = left; R = right; MCC = middle cingulate cortex; PCC = posterior cingulate cortex; STG = superior temporal gyrus; OFC = orbitofrontal cortex.

| Path | *t* | *p* | *p* (FDR) |
| --- | --- | --- | --- |
| *NCC vs. EP* |  |  |  |
| L.Putamen & L.MCC | -2.098 | 0.040 | 0.991 |
| R.Putamen & R.MCC | -2.139 | 0.037 | 0.991 |
| L.Putamen & R.MCC | -2.270 | 0.027 | 0.991 |
| *SA- vs. SA+* |  |  |  |
| L.OFC & R.Caudate | -2.014 | 0.049 | 0.992 |
| R.PCC & R.MCC | -2.330 | 0.023 | 0.992 |

## 4. Discussion

The present study is among the first to explore the neural correlates of NSSI in individuals with EP. Using GIMME to model resting-state effective connectivity among regions commonly implicated in self-injury studies, we found STG and insula driven connectivity to the PCC and striatum across the sample. NSSI was characterized by lagged PCC directed insula connectivity, contemporaneous right to left putamen connectivity, and stronger lagged than contemporaneous connectivity from the insula to the putamen. No circuits were specific to SA, but reduced PCC directed caudate connectivity was related to increased SIB more broadly. Disrupted neural circuitry associated with psychosis may further contribute to SIB risk by elevating right insula to right putamen and lagged right to left putamen connectivity. These findings suggest that PCC, insula, and putamen connectivity may be a marker for elevated SIB risk, and this overlap with psychosis-associated circuitry may contribute to the high prevalence of SIBs in psychosis.

Insula anomalies are consistently implicated in studies of SIBs. Many NSSI studies identify altered insula activity in both task (Bonenberger et al., 2015; R. C. Brown et al., 2017; Mayo et al., 2021; Minzenberg et al., 2016; Perini et al., 2019) and resting-state fMRI (Chen et al., 2023; Ho et al., 2021; Otto et al., 2023; Santamarina-Perez et al., 2019; Zhou et al., 2022). The insula contributes to interoception, self-referential processing, and emotion processing (Tisserand et al., 2023), all of which are implicated in SIB risk (Rogante et al., 2024; Seymour et al., 2016; L. Wang et al., 2021; Wolff, 2019). Notably, emotional reactivity and negative self-evaluation in NSSI is positively correlated with insula activation (Mayo et al., 2021; Perini et al., 2019). Our insula ROI comprises the middle insula, which responds to pain and interoceptive awareness (Kurth et al., 2010). Consistent with prior findings of altered insula response to pain in NSSI (Bonenberger et al., 2015), insula connectivity was a central feature of SIB history in our sample, suggesting that baseline disruptions in affective and interoceptive processing may contribute to self-harm in EP.

Lagged and contemporaneous insula directed to putamen, contemporaneous right to left putamen, and autoregressive putamen connectivity were also characteristic of NSSI. Striatal anomalies have been linked to both NSSI (Chen et al., 2023; Choi et al., 2024; Ho et al., 2021; X. Liu et al., 2025; Osuch et al., 2014; Poon et al., 2019; Santamarina-Perez et al., 2019; Sauder et al., 2016; K. Wang et al., 2022; Zhou et al., 2022) and STBs in psychosis (Minzenberg et al., 2015a, 2016). The putamen contributes to development of habitual behaviors and the caudate contributes to goal-directed behavior, both integral to reward processing (X. Liu et al., 2025). Given that NSSI may become a reinforced coping strategy over time, altered putamen connectivity may reflect changes in reward sensitivity and habit learning (R. T. Liu, 2017). Studies on NSSI find inconsistent results with reduced putamen activation during reward expectation (Sauder et al., 2016) and hyperactivity in response to rewarding stimuli (Osuch et al., 2014; Poon et al., 2019). Our findings of elevated contemporaneous but not lagged right to left putamen connectivity in NSSI could indicate enhanced within-reward-network processing, whereas stronger lagged than contemporaneous insula to putamen connectivity could represent disrupted interpretation of internal states, such as pain, as rewarding or reflect altered integration of interoceptive and reward signals. These results contribute to growing evidence of disrupted reward processing contributing to NSSI vulnerability.

Potentially catalyzing overall insula connectivity deviations, we found lagged left PCC to left insula connectivity specifically in participants with NSSI, and lack of right PCC to left caudate activity in participants with NSSI or SA. The PCC is involved in self-referential processing, rumination, and pain perception (Quevedo et al., 2016; Westlund Schreiner et al., 2023). NSSI is associated with more persistent rumination, with reduced PCC activation during rumination correlating with lifetime NSSI (Westlund Schreiner et al., 2023). Additionally, altered PCC activation is associated with reward processing and “relief” felt during self-inflicted pain (Osuch et al., 2014). This PCC to insula pathway may represent difficulties disengaging from rumination, fueling emotion dysregulation through perpetual negative emotional cycles. Reduced PCC to caudate connectivity related to SIB broadly may reflect disruptions in adaptive self-referential processing and reward anticipation (G. Liu et al., 2024; Osuch et al., 2014; Sauder et al., 2016). Deficits in adaptive self-referential processing and introspection may be driven by the PCC through disrupted emotion regulation and reward systems.

Several circuits characterized EP, NSSI, and SA. Contemporaneous and lagged insula to putamen connectivity was elevated in participants with EP, NSSI, or SA, suggesting shared neurobiological mechanisms underlying psychosis and self-injury. Existing studies on psychosis similarly report associations between insula and PCC anomalies and previous SA (Besteher et al., 2016; Canal-Rivero et al., 2025; Giakoumatos et al., 2013). Furthermore, increased contemporaneous and lagged left STG to left PCC connectivity was unique to psychosis and could exacerbate PCC-related vulnerabilities associated with NSSI. Consistent with this possibility, task-based fMRI studies in psychosis frequently identify aberrant cingulate activity related to STBs relative to patients with no STBs and healthy controls (Hoptman et al., 2024; Lee et al., 2015; Minzenberg et al., 2015b, 2016; H. Zhang et al., 2013). Overall, these results provide preliminary evidence that alterations in reward and emotion processing networks in psychosis may also increase vulnerability to SIBs.

### 4.1 Limitations and Future Directions

This study was limited by a relatively small sample and uneven distribution of SIB histories across EP and NCC groups, reducing our ability to distinguish neural components specific to NSSI or SA. Furthermore, we focused on lifetime histories and did not capture recency of SIBs. As a cross-sectional study, we cannot conclude that neural differences preceded or followed onset of SIB. Finally, although resting-state data provides valuable insight on baseline differences, these results can be difficult to relate to behavior due to a lack of task-based fMRI data assessing emotion or reward processing in the same participants.

Future studies should further explore the interaction between the neurobiological mechanisms of psychosis and NSSI. Longitudinal studies including clinical high-risk populations may clarify when these neural vulnerabilities emerge and how they relate to SIBs. Future work should also examine effective connectivity among additional cognitive control and pain-processing regions, including the PFC, anterior cingulate, inferior frontal gyrus, somatosensory cortex, and cuneus (**Supplemental Table 1b**). To preserve the stability of GIMME, only the most commonly significant regions were selected for this analysis, which resulted in limited representation.

## 5. Conclusion

Improving our understanding of neurobiological factors associated with SIBs in EP may support the development of interventions that effectively target suicidality in a highly at-risk population. NSSI provides important information about suicide risk but remains underrecognized in psychosis. Our findings emphasize the importance of screening for NSSI in addition to suicidal ideation or behaviors, as psychosis may confer a broader vulnerability to self-injury. This study uncovered patterns of PCC, insula, and striatal resting-state effective connectivity related to NSSI in EP, extending findings previously observed in other psychiatric disorders. Psychosis-related alterations involving the STG, PCC and insula may further increase vulnerability to SIB. Together, these findings underscore the importance of studying NSSI in EP and suggest that overlapping neural mechanisms may contribute to the elevated risk for both NSSI and suicide during this critical stage of illness.

## CRediT authorship contribution statement

**Cindy An**: Conceptualization, Data Curation, Formal analysis, Investigation, Methodology, Software, Visualization, Writing – original draft, Writing – review & editing. **Seema Dhaher**: Data Curation, Project Administration, Resources, Writing – review & editing. **Melissa Kilicoglu**: Data Curation, Project Administration, Resources, Writing – review & editing. **Jessica A. Turner**: Conceptualization, Methodology, Supervision, Writing – review & editing. **Mindy Westlund Schreiner**: Conceptualization, Methodology, Supervision, Writing – review & editing. **Aubrey Moe**: Conceptualization, Funding acquisition, Methodology, Project Administration, Resources, Supervision, Writing – review & editing.

## References

Aguilar, E. J., García-Martí, G., Martí-Bonmatí, L., Lull, J. J., Moratal, D., Escartí, M. J., Robles, M., González, J. C., Guillamón, M. I., & Sanjuán, J. (2008). Left orbitofrontal and superior temporal gyrus structural changes associated to suicidal behavior in patients with schizophrenia. Progress in Neuro-Psychopharmacology and Biological Psychiatry, 32(7), 1673–1676. 10.1016/j.pnpbp.2008.06.016

Athanassiou, M., Dumais, A., Iammatteo, V., De Benedictis, L., Dubreucq, J.-L., & Potvin, S. (2021). The processing of angry faces in schizophrenia patients with a history of suicide: An fMRI study examining brain activity and connectivity. Progress in Neuro-Psychopharmacology and Biological Psychiatry, 107, 110253. 10.1016/j.pnpbp.2021.110253

Auquier, P., Lançon, C., Rouillon, F., & Lader, M. (2007). Mortality in schizophrenia. Pharmacoepidemiology and Drug Safety, 16(12), 1308–1312. 10.1002/pds.1496

Benjamini, Y., & Hochberg, Y. (1995). Controlling the False Discovery Rate: A Practical and Powerful Approach to Multiple Testing. Journal of the Royal Statistical Society: Series B (Methodological), 57(1), 289–300. 10.1111/j.2517-6161.1995.tb02031.x

Besteher, B., Wagner, G., Koch, K., Schachtzabel, C., Reichenbach, J. R., Schlösser, R., Sauer, H., & Schultz, C. C. (2016). Pronounced prefronto-temporal cortical thinning in schizophrenia: Neuroanatomical correlate of suicidal behavior? Schizophrenia Research, 176(2), 151–157. 10.1016/j.schres.2016.08.010

Bonenberger, M., Plener, P. L., Groschwitz, R. C., Grön, G., & Abler, B. (2015). Differential neural processing of unpleasant haptic sensations in somatic and affective partitions of the insula in non-suicidal self-injury (NSSI). Psychiatry Research: Neuroimaging, 234(3), 298–304. 10.1016/j.pscychresns.2015.10.013

Brown, R. C., Plener, P. L., Groen, G., Neff, D., Bonenberger, M., & Abler, B. (2017). Differential Neural Processing of Social Exclusion and Inclusion in Adolescents with Non-Suicidal Self-Injury and Young Adults with Borderline Personality Disorder. Frontiers in Psychiatry, 8. 10.3389/fpsyt.2017.00267

Brown, S. (1997). Excess mortality of schizophrenia: A meta-analysis. The British Journal of Psychiatry, 171(6), 502–508. 10.1192/bjp.171.6.502

Buckner, R. L., Krienen, F. M., Castellanos, A., Diaz, J. C., & Yeo, B. T. T. (2011). The organization of the human cerebellum estimated by intrinsic functional connectivity. Journal of Neurophysiology, 106(5), 2322–2345. 10.1152/jn.00339.2011

Canal-Rivero, M., Tordesillas-Gutiérrez, D., Ruiz-Veguilla, M., Ortiz-García de la Foz, V., Cuevas-Esteban, J., Marco de Lucas, E., Vázquez-Bourgon, J., Ayesa-Arriola, R., & Crespo-Facorro, B. (2020). Brain grey matter abnormalities in first episode non-affective psychosis patients with suicidal behaviours: The role of neurocognitive functioning. Progress in Neuro-Psychopharmacology and Biological Psychiatry, 102, 109948. 10.1016/j.pnpbp.2020.109948

Canal-Rivero, M., Tordesillas-Gutiérrez, D., Ruiz-Veguilla, M., Ortiz-García de la Foz, V., Marco de Lucas, E., Romero-Garcia, R., Vázquez-Bourgon, J., Ayesa-Arriola, R., & Crespo-Facorro, B. (2025). Suicidal Behaviour Prior to First Episode Psychosis: Wider and More Widespread Grey-Matter Alterations. Archives of Suicide Research: Official Journal of the International Academy for Suicide Research, 1–15. 10.1080/13811118.2025.2454581

Cangur, S., & Ercan, I. (2015). Comparison of Model Fit Indices Used in Structural Equation Modeling Under Multivariate Normality. Journal of Modern Applied Statistical Methods, 14, 152–167. 10.56801/10.56801/v14.i.759

Chen, X., Chen, H., Liu, J., Tang, H., Zhou, J., Liu, P., Tian, Y., Wang, X., Lu, F., & Zhou, J. (2023). Functional connectivity alterations in reward-related circuits associated with non-suicidal self-injury behaviors in drug-naïve adolescents with depression. Journal of Psychiatric Research, 163, 270–277. 10.1016/j.jpsychires.2023.05.068

Choi, E. J., Vandewouw, M. M., Taylor, M. J., Stevenson, R. A., Arnold, P. D., Brian, J., Crosbie, J., Kelley, E., Liu, X., Jones, J., Lai, M.-C., Schachar, R. J., Lerch, J. P., & Anagnostou, E. (2024). Dorsal Striatal Functional Connectivity and Repetitive Behavior Dimensions in Children and Youths With Neurodevelopmental Disorders. Biological Psychiatry: Cognitive Neuroscience and Neuroimaging, 9(4), 387–397. 10.1016/j.bpsc.2023.10.014

Cooper, J., Kapur, N., Webb, R., Lawlor, M., Guthrie, E., Mackway-Jones, K., & Appleby, L. (2005). Suicide After Deliberate Self-Harm: A 4-Year Cohort Study. American Journal of Psychiatry, 162, 297–303. 10.1176/appi.ajp.162.2.297

Cullen, K. R., Westlund Schreiner, M., Klimes-Dougan, B., Eberly, L. E., LaRiviere, L. L., Lim, K. O., Camchong, J., & Mueller, B. A. (2020). Neural correlates of clinical improvement in response to N-acetylcysteine in adolescents with non-suicidal self-injury. Progress in Neuro-Psychopharmacology and Biological Psychiatry, 99, 109778. 10.1016/j.pnpbp.2019.109778

Demers, L. A., Westlund Schreiner, M., Hunt, R. H., Mueller, B. A., Klimes-Dougan, B., Thomas, K. M., & Cullen, K. R. (2019). Alexithymia is associated with neural reactivity to masked emotional faces in adolescents who self-harm. Journal of Affective Disorders, 249, 253–261. 10.1016/j.jad.2019.02.038

First, M. B., Williams, J. B., Karg, R. S., & Spitzer, R. L. (2015). Structured clinical interview for DSM-5 disorders. Clinician Version (SCID-5-CV).

Fischl, B., Salat, D. H., Busa, E., Albert, M., Dieterich, M., Haselgrove, C., van der Kouwe, A., Killiany, R., Kennedy, D., Klaveness, S., Montillo, A., Makris, N., Rosen, B., & Dale, A. M. (2002). Whole Brain Segmentation: Automated Labeling of Neuroanatomical Structures in the Human Brain. Neuron, 33(3), 341–355. 10.1016/S0896-6273(02)00569-X

Gates, K. M., & Molenaar, P. C. M. (2012). Group search algorithm recovers effective connectivity maps for individuals in homogeneous and heterogeneous samples. NeuroImage, 63(1), 310–319. 10.1016/j.neuroimage.2012.06.026

Giakoumatos, C. I., Tandon, N., Shah, J., Mathew, I. T., Brady, R. O., Clementz, B. A., Pearlson, G. D., Thaker, G. K., Tamminga, C. A., Sweeney, J. A., & Keshavan, M. S. (2013). Are Structural Brain Abnormalities Associated With Suicidal Behavior In Patients With Psychotic Disorders? Journal of Psychiatric Research, 47(10), 1389–1395. 10.1016/j.jpsychires.2013.06.011

Girgis, R. R., Basavaraju, R., France, J., Wall, M. M., Brucato, G., Lieberman, J. A., & Provenzano, F. A. (2021). An Exploratory Magnetic Resonance Imaging Study of Suicidal Ideation in Individuals at Clinical High-Risk for Psychosis. Psychiatry Research. Neuroimaging, 312, 111287. 10.1016/j.pscychresns.2021.111287

Harvey, S. B., Dean, K., Morgan, C., Walsh, E., Demjaha, A., Dazzan, P., Morgan, K., Lloyd, T., Fearon, P., Jones, P. B., & Murray, R. M. (2008). Self-harm in first-episode psychosis. The British Journal of Psychiatry, 192(3), 178–184. 10.1192/bjp.bp.107.037192

Hillary, F. G., Medaglia, J. D., Gates, K. M., Molenaar, P. C., & Good, D. C. (2014). Examining network dynamics after traumatic brain injury using the extended unified SEM approach. Brain Imaging and Behavior, 8(3), 435–445. 10.1007/s11682-012-9205-0

Ho, T. C., Walker, J. C., Teresi, G. I., Kulla, A., Kirshenbaum, J. S., Gifuni, A. J., Singh, M. K., & Gotlib, I. H. (2021). Default mode and salience network alterations in suicidal and non-suicidal self-injurious thoughts and behaviors in adolescents with depression. Translational Psychiatry, 11(1), 38. 10.1038/s41398-020-01103-x

Hoptman, M. J., Evans, K. T., Parincu, Z., Sparpana, A. M., Sullivan, E. F., Ahmed, A. O., & Iosifescu, D. V. (2024). Emotion-related impulsivity and suicidal ideation and behavior in schizophrenia spectrum disorder: A pilot fMRI study. Frontiers in Psychiatry, 15. 10.3389/fpsyt.2024.1408083

Huang, Q., Xiao, M., Ai, M., Chen, J., Wang, W., Hu, L., Cao, J., Wang, M., & Kuang, L. (2021). Disruption of Neural Activity and Functional Connectivity in Adolescents With Major Depressive Disorder Who Engage in Non-suicidal Self-Injury: A Resting-State fMRI Study. Frontiers in Psychiatry, 12. 10.3389/fpsyt.2021.571532

Keane, B. P., Abrham, Y. T., Hearne, L. J., Bi, H., & Hu, B. (2024). Increased whole-brain functional heterogeneity in psychosis during rest and task. NeuroImage: Clinical, 43, 103630. 10.1016/j.nicl.2024.103630

Kurdyak, P., Mallia, E., de Oliveira, C., Carvalho, A. F., Kozloff, N., Zaheer, J., Tempelaar, W. M., Anderson, K. K., Correll, C. U., & Voineskos, A. N. (2021). Mortality After the First Diagnosis of Schizophrenia-Spectrum Disorders: A Population-based Retrospective Cohort Study. Schizophrenia Bulletin, 47(3), 864–874. 10.1093/schbul/sbaa180

Kurth, F., Eickhoff, S. B., Schleicher, A., Hoemke, L., Zilles, K., & Amunts, K. (2010). Cytoarchitecture and Probabilistic Maps of the Human Posterior Insular Cortex. *Cerebral Cortex (New York*, NY), 20(6), 1448–1461. 10.1093/cercor/bhp208

Lane, S., Gates, K. M., Fisher, Z., Arizmendi, C., & Molenaar, P. (2025). gimme: Group Iterative Multiple Model Estimation (Version R package version 0.8.3) [Computer software]. https://CRAN.R-project.org/package=gimme

Laursen, T. M., Nordentoft, M., & Mortensen, P. B. (2014). Excess Early Mortality in Schizophrenia. Annual Review of Clinical Psychology, 10(Volume 10, 2014), 425–448. 10.1146/annurev-clinpsy-032813-153657

Lee, K.-H., Pluck, G., Lekka, N., Horton, A., Wilkinson, I. D., & Woodruff, P. W. R. (2015). Self-harm in schizophrenia is associated with dorsolateral prefrontal and posterior cingulate activity. Progress in Neuro-Psychopharmacology and Biological Psychiatry, 61, 18–23. 10.1016/j.pnpbp.2015.03.005

Li, M., Xiao, Y., Ge, Y., Yan, H., Li, X., Yue, W., & Yan, H. (2026). Functional heterogeneity in non-suicidal self-injury across psychiatric disorders: Neural and psychosocial correlates. Translational Psychiatry, 16(1), 60. 10.1038/s41398-025-03802-9

Liao, K., Yu, R., Chen, Y., Chen, X., Wu, X., Huang, X., & Liu, N. (2024). Alterations of regional brain activity and corresponding brain circuits in drug-naïve adolescents with nonsuicidal self-injury. Scientific Reports, 14(1), 24997. 10.1038/s41598-024-75714-5

Liu, G., Hao, G., Das, N., Ranatunga, J., Schneider, C., Yang, L., & Quevedo, K. (2024). Self-compassion, self-referential caudate circuitry, and adolescent suicide ideation. Translational Psychiatry, 14, 334. 10.1038/s41398-024-03037-0

Liu, R. T. (2017). Characterizing the course of non-suicidal self-injury: A cognitive neuroscience perspective. Neuroscience and Biobehavioral Reviews, 80, 159–165. 10.1016/j.neubiorev.2017.05.026

Liu, X., Zhang, Y., Chen, J., Xie, M., Pan, L., Hommel, B., Yang, Y., Zhu, X., Wang, K., & Zhang, W. (2025). Altered brain structure and function correlate with non-suicidal self-injury in children and adolescents with transdiagnostic psychiatric disorders. Journal of Psychiatric Research, 184, 17–26. 10.1016/j.jpsychires.2025.02.051

Lorentzen, E. A., Mors, O., & Kjær, J. N. (2022). The Prevalence of Self-injurious Behavior in Patients With Schizophrenia Spectrum Disorders: A Systematic Review and Meta-analysis. Schizophrenia Bulletin Open, 3(1), sgac069. 10.1093/schizbullopen/sgac069

Mayo, L. M., Perini, I., Gustafsson, P. A., Hamilton, J. P., Kämpe, R., Heilig, M., & Zetterqvist, M. (2021). Psychophysiological and Neural Support for Enhanced Emotional Reactivity in Female Adolescents With Nonsuicidal Self-injury. Biological Psychiatry: Cognitive Neuroscience and Neuroimaging, 6(7), 682–691. 10.1016/j.bpsc.2020.11.004

Meltzer, H. Y. (2001). Treatment of suicidality in schizophrenia. Annals of the New York Academy of Sciences, 932, 44–58; discussion 58-60. 10.1111/j.1749-6632.2001.tb05797.x

Minzenberg, M. J., Lesh, T. A., Niendam, T. A., Cheng, Y., & Carter, C. S. (2016). Conflict-Related Anterior Cingulate Functional Connectivity Is Associated With Past Suicidal Ideation and Behavior in Recent-Onset Psychotic Major Mood Disorders. The Journal of Neuropsychiatry and Clinical Neurosciences, 28(4), 299–305. 10.1176/appi.neuropsych.15120422

Minzenberg, M. J., Lesh, T. A., Niendam, T. A., Yoon, J. H., Cheng, Y., Rhoades, R. N., & Carter, C. S. (2015a). Control-related frontal-striatal function is associated with past suicidal ideation and behavior in patients with recent-onset psychotic major mood disorders. Journal of Affective Disorders, 188, 202–209. 10.1016/j.jad.2015.08.049

Minzenberg, M. J., Lesh, T., Niendam, T., Yoon, J. H., Cheng, Y., Rhoades, R. N., & Carter, C. S. (2015b). Frontal Motor Cortex Activity During Reactive Control Is Associated With Past Suicidal Behavior in Recent-Onset Schizophrenia. Crisis, 36(5), 363–370. 10.1027/0227-5910/a000335

Moe, A. M., Llamocca, E., Wastler, H. M., Steelesmith, D. L., Brock, G., Bridge, J. A., & Fontanella, C. A. (2022). Risk Factors for Deliberate Self-harm and Suicide Among Adolescents and Young Adults With First-Episode Psychosis. Schizophrenia Bulletin, 48(2), 414–424. 10.1093/schbul/sbab123

Mortensen, P. B., & Juel, K. (1990). Mortality and causes of death in schizophrenic patients in Denmark. Acta Psychiatrica Scandinavica, 81(4), 372–377. 10.1111/j.1600-0447.1990.tb05466.x

Mortensen, P. B., & Juel, K. (1993). Mortality and Causes of Death in First Admitted Schizophrenic Patients. The British Journal of Psychiatry, 163(2), 183–189. 10.1192/bjp.163.2.183

Nanda, P., Tandon, N., Mathew, I. T., Padmanabhan, J. L., Clementz, B. A., Pearlson, G. D., Sweeney, J. A., Tamminga, C. A., & Keshavan, M. S. (2016). Impulsivity across the psychosis spectrum: Correlates of cortical volume, suicidal history, and social and global function. Schizophrenia Research, 170(1), 80–86. 10.1016/j.schres.2015.11.030

Nieto, E., Vieta, E., Gastó, C., Vallejo, J., & Cirera, E. (1992). Suicide attempts of high medical seriousness in schizophrenic patients. Comprehensive Psychiatry, 33(6), 384–387. 10.1016/0010-440x(92)90060-4

Nock, M. K., Holmberg, E. B., Photos, V. I., & Michel, B. D. (2007). Self-Injurious Thoughts and Behaviors Interview: Development, reliability, and validity in an adolescent sample. Psychological Assessment, 19(3), 309–317. 10.1037/1040-3590.19.3.309

Osuch, E., Ford, K., Wrath, A., Bartha, R., & Neufeld, R. (2014). Functional MRI of pain application in youth who engaged in repetitive non-suicidal self-injury vs. Psychiatric controls. Psychiatry Research: Neuroimaging, 223(2), 104–112. 10.1016/j.pscychresns.2014.05.003

Otto, A., Jarvers, I., Kandsperger, S., Reichl, C., Ando, A., Koenig, J., Kaess, M., & Brunner, R. (2023). Stress-induced alterations in resting-state functional connectivity among adolescents with non-suicidal self-injury. Journal of Affective Disorders, 339, 162–171. 10.1016/j.jad.2023.07.032

Palmer, B. A., Pankratz, V. S., & Bostwick, J. M. (2005). The Lifetime Risk of Suicide in Schizophrenia: A Reexamination. Archives of General Psychiatry, 62(3), 247–253. 10.1001/archpsyc.62.3.247

Perini, I., Gustafsson, P. A., Hamilton, J. P., Kämpe, R., Mayo, L. M., Heilig, M., & Zetterqvist, M. (2019). Brain-based Classification of Negative Social Bias in Adolescents With Nonsuicidal Self-injury: Findings From Simulated Online Social Interaction. EClinicalMedicine, 13, 81–90. 10.1016/j.eclinm.2019.06.016

Plener, P. L., Bubalo, N., Fladung, A. K., Ludolph, A. G., & Lulé, D. (2012). Prone to excitement: Adolescent females with non-suicidal self-injury (NSSI) show altered cortical pattern to emotional and NSS-related material. Psychiatry Research: Neuroimaging, 203(2), 146–152. 10.1016/j.pscychresns.2011.12.012

Poon, J. A., Thompson, J. C., Forbes, E. E., & Chaplin, T. M. (2019). Adolescents’ Reward-related Neural Activation: Links to Thoughts of Nonsuicidal Self-Injury. Suicide and Life-Threatening Behavior, 49(1), 76–89. 10.1111/sltb.12418

Potvin, S., Tikàsz, A., Richard-Devantoy, S., Lungu, O., & Dumais, A. (2018). History of Suicide Attempt Is Associated with Reduced Medial Prefrontal Cortex Activity during Emotional Decision-Making among Men with Schizophrenia: An Exploratory fMRI Study. Schizophrenia Research and Treatment, 2018, 9898654. 10.1155/2018/9898654

Quevedo, K., Martin, J., Scott, H., Smyda, G., & Pfeifer, J. H. (2016). The neurobiology of self-knowledge in depressed and self-injurious youth. Psychiatry Research: Neuroimaging, 254, 145–155. 10.1016/j.pscychresns.2016.06.015

R Core Team. (2024). R: A Language and Environment for Statistical Computing [Computer software]. R Foundation for Statistical Computing. https://www.R-project.org/

Reitz, S., Kluetsch, R., Niedtfeld, I., Knorz, T., Lis, S., Paret, C., Kirsch, P., Meyer-Lindenberg, A., Treede, R.-D., Baumgärtner, U., Bohus, M., & Schmahl, C. (2015). Incision and stress regulation in borderline personality disorder: Neurobiological mechanisms of self-injurious behaviour. The British Journal of Psychiatry, 207(2), 165–172. 10.1192/bjp.bp.114.153379

Robinson, J., Harris, M. G., Harrigan, S. M., Henry, L. P., Farrelly, S., Prosser, A., Schwartz, O., Jackson, H., & McGorry, P. D. (2010). Suicide attempt in first-episode psychosis: A 7.4 year follow-up study. Schizophrenia Research, 116(1), 1–8. 10.1016/j.schres.2009.10.009

Rogante, E., Cifrodelli, M., Sarubbi, S., Costanza, A., Erbuto, D., Berardelli, I., & Pompili, M. (2024). The Role of Emotion Dysregulation in Understanding Suicide Risk: A Systematic Review of the Literature. Healthcare, 12(2), 169. 10.3390/healthcare12020169

Rüsch, N., Spoletini, I., Wilke, M., Martinotti, G., Bria, P., Trequattrini, A., Bonaviri, G., Caltagirone, C., & Spalletta, G. (2008). Inferior frontal white matter volume and suicidality in schizophrenia. Psychiatry Research: Neuroimaging, 164(3), 206–214. 10.1016/j.pscychresns.2007.12.011

Santamarina-Perez, P., Romero, S., Mendez, I., Leslie, S. M., Packer, M. M., Sugranyes, G., Picado, M., Font, E., Moreno, E., Martinez, E., Morer, A., Romero, M., & Singh, M. K. (2019). Fronto-Limbic Connectivity as a Predictor of Improvement in Nonsuicidal Self-Injury in Adolescents Following Psychotherapy. Journal of Child and Adolescent Psychopharmacology, 29(6), 456–465. 10.1089/cap.2018.0152

Sauder, C. L., Derbidge, C. M., & Beauchaine, T. P. (2016). Neural responses to monetary incentives among self-injuring adolescent girls. Development and Psychopathology, 28(1), 277–291. 10.1017/S0954579415000449

Schaefer, A., Kong, R., Gordon, E. M., Laumann, T. O., Zuo, X.-N., Holmes, A. J., Eickhoff, S. B., & Yeo, B. T. T. (2018). Local-Global Parcellation of the Human Cerebral Cortex from Intrinsic Functional Connectivity MRI. Cerebral Cortex, 28(9), 3095–3114. 10.1093/cercor/bhx179

Schmaal, L., van Harmelen, A.-L., Chatzi, V., Lippard, E. T. C., Toenders, Y. J., Averill, L. A., Mazure, C. M., & Blumberg, H. P. (2020). Imaging suicidal thoughts and behaviors: A comprehensive review of 2 decades of neuroimaging studies. Molecular Psychiatry, 25(2), 408–427. 10.1038/s41380-019-0587-x

Seymour, K. E., Jones, R. N., Cushman, G. K., Galvan, T., Puzia, M. E., Kim, K. L., Spirito, A., & Dickstein, D. P. (2016). Emotional face recognition in adolescent suicide attempters and adolescents engaging in non-suicidal self-injury. European Child & Adolescent Psychiatry, 25(3), 247–259. 10.1007/s00787-015-0733-1

Spoletini, I., Piras, F., Fagioli, S., Rubino, I. A., Martinotti, G., Siracusano, A., Caltagirone, C., & Spalletta, G. (2011). Suicidal attempts and increased right amygdala volume in schizophrenia. Schizophrenia Research, 125(1), 30–40. 10.1016/j.schres.2010.08.023

Symonds, C. S., Taylor, S., Tippins, V., & Turkington, D. (2006). Violent Self-Harm in Schizophrenia. Suicide and Life-Threatening Behavior, 36(1), 44–49. 10.1521/suli.2006.36.1.44

Thomas Yeo, B. T., Krienen, F. M., Sepulcre, J., Sabuncu, M. R., Lashkari, D., Hollinshead, M., Roffman, J. L., Smoller, J. W., Zöllei, L., Polimeni, J. R., Fischl, B., Liu, H., & Buckner, R. L. (2011). The organization of the human cerebral cortex estimated by intrinsic functional connectivity. Journal of Neurophysiology, 106(3), 1125–1165. 10.1152/jn.00338.2011

Tisserand, A., Philippi, N., Botzung, A., & Blanc, F. (2023). Me, Myself and My Insula: An Oasis in the Forefront of Self-Consciousness. Biology, 12(4), 599. 10.3390/biology12040599

Vega, D., Ripollés, P., Soto, À., Torrubia, R., Ribas, J., Monreal, J. A., Pascual, J. C., Salvador, R., Pomarol-Clotet, E., Rodríguez-Fornells, A., & Marco-Pallarés, J. (2018). Orbitofrontal overactivation in reward processing in borderline personality disorder: The role of non-suicidal self-injury. Brain Imaging and Behavior, 12(1), 217–228. 10.1007/s11682-017-9687-x

Waller, L., Erk, S., Pozzi, E., Toenders, Y. J., Haswell, C. C., Büttner, M., Thompson, P. M., Schmaal, L., Morey, R. A., Walter, H., & Veer, I. M. (2022). ENIGMA HALFpipe: Interactive, reproducible, and efficient analysis for resting-state and task-based fMRI data. Human Brain Mapping, 43(9), 2727–2742. 10.1002/hbm.25829

Wang, K., He, Q., Zhu, X., Hu, Y., Yao, Y., Hommel, B., Beste, C., Liu, J., Yang, Y., & Zhang, W. (2022). Smaller putamen volumes are associated with greater problems in external emotional regulation in depressed adolescents with nonsuicidal self-injury. Journal of Psychiatric Research, 155, 338–346. 10.1016/j.jpsychires.2022.09.014

Wang, L., Cui, Q., Liu, J., & Zou, H. (2021). Emotion Reactivity and Suicide Risk in Patients With Depression: The Mediating Role of Non-Suicidal Self-Injury and Moderating Role of Childhood Neglect. Frontiers in Psychiatry, 12, 707181. 10.3389/fpsyt.2021.707181

Waugh, A. C. (1986). Autocastration and biblical delusions in schizophrenia. The British Journal of Psychiatry: The Journal of Mental Science, 149, 656–658. 10.1192/bjp.149.5.656

Westlund Schreiner, M., Klimes-Dougan, B., Mueller, B. A., Eberly, L. E., Reigstad, K. M., Carstedt, P. A., Thomas, K. M., Hunt, R. H., Lim, K. O., & Cullen, K. R. (2017). Multi-modal neuroimaging of adolescents with non-suicidal self-injury: Amygdala functional connectivity. Journal of Affective Disorders, 221, 47–55. 10.1016/j.jad.2017.06.004

Westlund Schreiner, M., Roberts, H., Dillahunt, A. K., Farstead, B., Feldman, D., Thomas, L., Jacobs, R. H., Bessette, K. L., Welsh, R. C., Watkins, E. R., Langenecker, S. A., & Crowell, S. E. (2023). Negative association between non-suicidal self-injury in adolescents and default mode network activation during the distraction blocks of a rumination task. Suicide and Life-Threatening Behavior, 53(3), 510–521. 10.1111/sltb.12960

Wilkinson, G. S., & Robertson, G. J. (1993). Wide range achievement test 4. Journal of Clinical and Experimental Neuropsychology.

Wolff, J. (2019). Emotion Dysregulation and Non-Suicidal Self-Injury: A Systematic Review and Meta-Analysis. European Psychiatry: The Journal of the Association of European Psychiatrists, 59, 25–36. 10.1016/j.eurpsy.2019.03.004

Yin, Y., Tong, J., Huang, J., Wang, L., Tian, B., Chen, S., Tan, S., Wang, Z., Yu, T., Li, Y., Tong, Y., Fan, F., Kochunov, P., Hong, L. E., & Tan, Y. (2023). History of suicide attempt associated with amygdala and hippocampus changes among individuals with schizophrenia. European Archives of Psychiatry and Clinical Neuroscience, 273(4), 921–930. 10.1007/s00406-023-01554-5

Zaheer, J., Olfson, M., Mallia, E., Lam, J. S. H., de Oliveira, C., Rudoler, D., Carvalho, A. F., Jacob, B. J., Juda, A., & Kurdyak, P. (2020). Predictors of suicide at time of diagnosis in schizophrenia spectrum disorder: A 20-year total population study in Ontario, Canada. Schizophrenia Research, 222, 382–388. 10.1016/j.schres.2020.04.025

Zhang, H., Wei, X., Tao, H., Mwansisya, T. E., Pu, W., He, Z., Hu, A., Xu, L., Liu, Z., Shan, B., & Xue, Z. (2013). Opposite Effective Connectivity in the Posterior Cingulate and Medial Prefrontal Cortex between First-Episode Schizophrenic Patients with Suicide Risk and Healthy Controls. PLOS ONE, 8(5), e63477. 10.1371/journal.pone.0063477

Zhou, Y., Yu, R., Ai, M., Cao, J., Li, X., Hong, S., Huang, Q., Dai, L., Wang, L., Zhao, L., Zhang, Q., Shi, L., & Kuang, L. (2022). A Resting State Functional Magnetic Resonance Imaging Study of Unmedicated Adolescents With Non-suicidal Self-Injury Behaviors: Evidence From the Amplitude of Low-Frequency Fluctuation and Regional Homogeneity Indicator. Frontiers in Psychiatry, 13. 10.3389/fpsyt.2022.925672

