## Supplement for "Anomalous Emotion Regulation & Reward Network Connectivity Underlying Suicidal & Non-Suicidal Self-Injury in Early Psychosis"

*1. ROI Determination*

We conducted a general review to identify significant regions of interest consistently identified in NSSI in adolescents and suicidality in psychosis studies. Articles were identified using searches through PubMed and PsycINFO. The bibliographies for the search-identified articles were reviewed so articles that matched criteria but were not previously identified could be included.

Non-suicidal self-injury:

- PubMed: "Self-Injurious Behavior"[Mesh] AND "Functional Neuroimaging"[Mesh] AND "Adolescent"[Mesh] AND “Self-inflicted injury”
- PsycINFO: (self-injury or self-harm or self-mutilation or self-mutilating or self-injurious) OR (self-injurious behavior or self injury or self harm)

Suicidality and psychosis:

- Automatically included all the papers from the Girgis, 2020 review.
- PubMed: "Suicide"[Mesh] AND "Functional Neuroimaging"[Mesh] AND "Psychotic Disorders"[Mesh]
- PubMed: "Self-Injurious Behavior"[Mesh] AND "Functional Neuroimaging"[Mesh] AND "Psychotic Disorders"[Mesh]
- PsycINFO: (suicide or suicidal ideation or suicidality or suicide attempts) AND psychosis AND (mri or magnetic resonance imaging or mri scan or mr)
- PsycINFO: (self harm or non-suicidal self-injury) AND psychosis AND (mri or magnetic resonance imaging or mri scan or mr)

This search identified 37 peer-reviewed articles on MRI studies about NSSI in adolescents and 21 articles about the neurobiology of suicidality in psychosis. No articles were found about the neurobiology of NSSI in psychosis. All significant areas identified within the cingulate and striatum were cumulatively counted. The most commonly occurring significant regions were chosen for the GIMME analysis (**Supplemental Table 1a & b**). Because of naming variability for regions within the cingulate, MNI coordinates for the posterior, middle, and anterior cingulate were plotted together and overlayed on the Schaefer atlas to determine the primary representative clusters. The medial frontal gyrus was a commonly significant region but was not selected due to extreme variation in significant coordinates between studies and limited region selection for the following effective connectivity analysis.

| **Publication** | **Category** | **Imaging Measure** | **MCC** | **PCC** | **Caudate** | **Putamen** | **STG** | **OFC** | **INS** |
| --- | --- | --- | --- | --- | --- | --- | --- | --- | --- |
| (Aguilar et al., 2008) | Psychosis | Gray matter voxel-based morphometry | 0 | 0 | 0 | 0 | -1  (−40, −31, 11) | -1  (−9, 39, −21) | 0 |
| (Beauchaine et al., 2019) | NSSI | Gray matter voxel-based morphometry | 0 | 0 | 0 | 0 | 0 | 0 | -1  (−38, −10, −14),  (33, 33, 6),  (34, −18, 7) |
| (Canal-Rivero et al., 2020) | Psychosis | Gray matter voxel-based morphometry | -1 (-9, -27, 34) | 0 | 0 | 0 | -1  (27, 20, −32),  (38, 12, −45) | 0 | 0 |
| (Canal-Rivero et al., 2025) | Psychosis | Gray matter voxel-based morphometry | 0 | -1  (−4, −60, 26) | 0 | 0 | -1  (−54, −40, 20) | 0 | 0 |
| (Liu et al., 2025) | NSSI | Gray matter voxel-based morphometry and resting-state fMRI connectivity | 0 | 0 | 0 | -1  (-31.5, -3, 6), (31.5, -3, 9) | -1  (-45, -36, 0), (48, -27, 3), (-51, -39, 6) | 0 | -1  (-31.5, -3, 6)  (31.5, -3, 9) |
| (Whittle et al., 2009) | NSSI | Gray matter volumetry and functional | 0 | 0 | 0 | 0 | 0 | 0 | 0 |
| (Takahashi et al., 2009) | NSSI | Gray matter volumetry | 0 | 0 | 0 | 0 | 0 | 0 | 0 |
| (Takahashi et al., 2010) | NSSI | Gray matter volumetry | 0 | 0 | 0 | 0 | 0 | 0 | 0 |
| (Spoletini et al., 2011) | Psychosis | Gray matter volumetry | 0 | 0 | 0 | 0 | 0 | 0 | 0 |
| (Giakoumatos et al., 2013) | Psychosis | Gray matter volumetry | 0 | 0 | 0 | 0 | -1 | -1 | -1 |
| (Nanda et al., 2016) | Psychosis | Gray matter volumetry | 0 | 0 | 0 | 0 | 0 | -1 | 0 |
| (Ando et al., 2018) | NSSI | Gray matter volumetry | 0 | 0 | 0 | 0 | 0 | 0 | -1 |
| (Yin et al., 2023) | Psychosis | Gray matter volumetry | 0 | 0 | 0 | 0 | 0 | 0 | 0 |
| (Wang et al., 2022) | NSSI | Gray matter volumetry and cortical thickness | 0 | 0 | 0 | -1 | 0 | 0 | 0 |
| (Besteher et al., 2016) | Psychosis | Gray matter cortical thickness | 0 | 0 | 0 | 0 | -1  (42, 21, - 35) | 0 | -1  (42, 21, - 35) |
| (Girgis et al., 2021) | Psychosis | Gray matter cortical thickness | 0 | 0 | 0 | 0 | 1 | 0 | 1 |
| (Huber et al., 2025) | NSSI | Gray matter cortical thickness and volumetry | 0 | 0 | 0 | 0 | 0 | 0 | 0 |
| (Rüsch et al., 2008) | Psychosis | Gray matter voxel-based morphometry and white matter volumetry | 0 | 0 | 0 | 0 | 0 | 1  (-31, 25, -7) | 0 |
| (Duerden et al., 2014) | NSSI | Gray matter cortical thickness and white matter FA & MD | 0 | 0 | 0 | 0 | 0 | 0 | 0 |
| (S.-J. Lee et al., 2016) | Psychosis | White matter FA & MD | 1 | 0 | 1 | 1 | 1 | 0 | 1 |
| (Long et al., 2018) | Psychosis | White matter FA & MD | 1 | 0 | 0 | 0 | 1 | 0 | 1 |
| (Patel et al., 2023) | NSSI | Gray matter cortical surface area | 0 | 0 | 0 | 0 | 0 | 0 | -1  (-35.2 5.0 14.0) |
| (Plener et al., 2012) | NSSI | fMRI task activation | 1  (-2, 7, 26) | 0 | 0 | 0 | 1  (-60, 44, 21) | 1  (-10, 46, -12) | 0 |
| (Minzenberg et al., 2014) | Psychosis | fMRI task activation | 0 | 0 | 0 | 0 | 0 | 0 | 0 |
| (Bonenberger et al., 2015) | NSSI | fMRI task activation | 0 | 0 | 0 | 0 | 0 | 0 | -1  (-39, -1, -6), (-34, 14, -3), (37, 14, -1) |
| (K.-H. Lee et al., 2015) | Psychosis | fMRI task activation | 0 | 1  (2, -20, 27) | 0 | 0 | 0 | 0 | 0 |
| (Minzenberg et al., 2015a) | Psychosis | fMRI task activation | 0 | -1  (-2, -66, 42), (0, -56, 44), (-2, -72, 34) | 1  (8, 2, 4) | 1  (8, 2, 4) | 0 | 1  (12, 24, 4) | 1  (8, 2, 4), (44, 0, 2) |
| (Minzenberg et al., 2015b) | Psychosis | fMRI task activation | 0 | 0 | 0 | 0 | 0 | 0 | 0 |
| (Groschwitz et al., 2016) | NSSI | fMRI task activation | 0 | 0 | 0 | 0 | 0 | 0 | 0 |
| (Sauder et al., 2016) | NSSI | fMRI task activation | 0 | 0 | 0 | -1  (33, -12, 9), (-27, -18, 6) | 0 | -1  (33, 48, -3), (-3, 63, -6) | 0 |
| (Quevedo et al., 2016) | NSSI | fMRI task activation | 1  (12, -40, 48) | 1  (12, -40, 48) | 0 | 0 | 0 | 0 | 0 |
| (Brown et al., 2017) | NSSI | fMRI task activation | 0 | 0 | 0 | 1  (-18, 16, -4) | 0 | 0 | -1  (-28, 16, 16) |
| (Potvin et al., 2018) | Psychosis | fMRI task activation | 0 | 0 | 0 | 0 | 1  (18, -64, -29) | 0 | 0 |
| (Demers et al., 2019) | NSSI | fMRI task activation | 0 | 0 | 0 | 0 | 0 | 0 | 0 |
| (Poon et al., 2019) | NSSI | fMRI task activation | 0 | 0 | 0 | 1 | 0 | 0 | 0 |
| (Mayo et al., 2021) | NSSI | fMRI task activation | 0 | 0 | 0 | 0 | 0 | 0 | 1 |
| (Malejko et al., 2022) | NSSI | fMRI task activation | 0 | 0 | 0 | 0 | 0 | 0 | 0 |
| (Westlund Schreiner et al., 2023) | NSSI | fMRI task activation | 0 | 1  (-6, -62, 16) | 0 | 0 | 0 | 0 | 0 |
| (Hoptman et al., 2024) | Psychosis | fMRI task activation | 1  (3, -23, 41) | 0 | 0 | 0 | 1  (59, -36, 11) | 0 | 0 |
| (Athanassiou et al., 2021) | Psychosis | fMRI task activation & connectivity | 1  (-4, -22, 40), (-4, 8, 30) | 0 | 0 | 0 | 0 | 0 | 0 |
| (Osuch et al., 2014) | NSSI | fMRI task seed-based connectivity | 0 | 1  (18, -52, 32), (-18, -76, 30) | 1  (12, -2, 10) | 0 | 1  (38, 6, -26) | -1  (24, 42, -20) | 0 |
| (Minzenberg et al., 2016) | Psychosis | fMRI task seed-based connectivity | -1  (6, 4, 40), (-8, -2, 44) | -1  (6, 4, 40), (-8, -2, 44) | 0 | -1  (22, 16, -4), (-30, 6, -2) | 1  (-48, -60, 30) | -1  (-26, 32, -10) | -1  (-38, 18, -4) |
| (L. Lin et al., 2024) | NSSI | fMRI task seed-based connectivity | 0 | 0 | 0 | 0 | 0 | 1  (-6, 40, -10) | 0 |
| (Santana-Gonzalez et al., 2025) | NSSI | fMRI task seed-based connectivity | 0 | 1  (14, -50, -2) | 0 | 0 | 1  (-63, -26, 45) | 0 | 0 |
| (H. Zhang et al., 2013) | Psychosis | fMRI task effective connectivity | 0 | -1  (0, -49, 28) | 0 | 0 | 0 | 0 | 0 |
| (Perini et al., 2019) | NSSI | fMRI task MVPA classification | 0 | 1  (8, -58, 37) | 0 | 0 | 1  (-47, -66, 15) | 1  (24, 16, -19), (-25, 22, -18), (-22, 12, -19), (9, 24 ,-17) | 1  (-45, -38, 19), (36, -22, 8) |
| (Westlund Schreiner et al., 2017) | NSSI | fMRI task and resting state seed-based connectivity | 0 | 0 | 0 | 0 | -1  (-58, -26, -8) | 0 | 0 |
| (Santamarina-Perez et al., 2019) | NSSI | fMRI resting state seed-based connectivity | 0 | 0 | 1 | 1 | 0 | 0 | -1  (56, -26, 14), (-34, -8, 12) |
| (Cullen et al., 2020) | NSSI | fMRI resting state seed-based connectivity | 0 | 0 | 0 | 0 | 0 | 0 | 0 |
| (Otto et al., 2023) | NSSI | fMRI resting state seed-based connectivity | 0 | 0 | 0 | 0 | 0 | 0 | 1 |
| (Choi et al., 2024) | NSSI | fMRI resting state seed-based connectivity | 0 | 0 | -1 | -1 | 0 | 0 | 0 |
| (Huang et al., 2021) | NSSI | fMRI resting state ALFF and seed-based connectivity | 1  (12, -24, 39) | -1  (-9, -39, 27) | 0 | 0 | -1  (63, -33, 3) | -1  (0, 63, 18) | 0 |
| (Liao et al., 2024) | NSSI | fMRI resting state ALFF and seed-based connectivity | -1  (9, 30, 32) | 0 | 0 | 0 | -1  (-48, 0, 0) | 0 | 0 |
| (Zhou et al., 2022) | NSSI | fMRI resting state ALFF, fALFF, and ReHo | 1  (-9, -42, 39), (12, -12, 42) | 1  (-3, -39, 24) | -1  (12, -3, 18) | 0 | 1  (45, 0, -9) | 0 | 1  (36, 15, -9), (45, -3, 3) |
| (X. Lin et al., 2025) | NSSI | fMRI resting state ALFF and resting state functional connectivity | 0 | 0 | 0 | 0 | 0 | 0 | 0 |
| (Chen et al., 2023) | NSSI | fMRI resting state functional connectivity | -1 | -1 | -1 | -1 | 0 | 0 | -1  (33, 5, 14), (33, 6, 14) |
| (Ho et al., 2021) | NSSI | fMRI resting state ICA and seed-based connectivity | 0 | -1 | -1 | -1 | 0 | 0 | -1 |
| (J. Zhang et al., 2025) | NSSI | fMRI resting state ICA and voxel-wise connectivity | 1  (-3, -33, 45) | 0 | 0 | 0 | 0 | 0 | 0 |
| **Psychosis Total** | | | 6 (29%) | 5 (24%) | 2 (10%) | 3 (14%) | 11 (52%) | 6 (29%) | 7 (33%) |
| **NSSI Total** | | | 7 (19%) | 9 (24%) | 6 (16%) | 9 (24%) | 9 (24%) | 6 (16%) | 13 (35%) |
| **Overall Total** | | | **13 (22%)** | **14 (24%)** | **8 (14%)** | **12 (21%)** | **20 (34%)** | **12 (21%)** | **20 (34%)** |

**Supplemental Table 1a.** Significant ROIs present in literature from adolescents with NSSI and suicidality in psychosis that were selected for GIMME analysis. Coordinates in MNI space were listed if they were available. +1 = present and increased; -1 = present and decreased; 0 = not present. FA = fractional anisotropy; MD = mean diffusivity; NSSI = non-suicidal self-injury; MCC = middle cingulate cortex; PCC = posterior cingulate cortex; STG = superior temporal gyrus; OFC = orbitofrontal cortex.

| **Publication** | **Category** | **Imaging Measure** | **Left Hemisphere** | **Right Hemisphere** |
| --- | --- | --- | --- | --- |
| (Aguilar et al., 2008) | Psychosis | Gray matter voxel-based morphometry |  |  |
| (Beauchaine et al., 2019) | NSSI | Gray matter voxel-based morphometry |  | IFG (33, 33, 6) |
| (Canal-Rivero et al., 2020) | Psychosis | Gray matter voxel-based morphometry | MFG (0, 50, 20)  Middle frontal gyrus (28, 42, 24)  Precentral gyrus (-34, -10, 50), (-44, -8, 38), (-48, -3, 30), (-62, 0, 15), (-27, -9, 64)  Subcallosal gyrus (-4, 6, -14)  Cuneus (-15, -94, 2) | IFG (44, 14, 21)  MFG (12, 30, 36), (4, 45, 40), (12, 38, 30), (2, 50, 28), (28, 42, 24)  Middle FG (-24, -4, 45)  Precentral Gyrus (14, -40, 45)  Precuneus (14, -40, 45)  Uncus (22, 9, -28), (10, -4, -26)  ACC (9, 22, -8)  Paracentral lobule (3, -30, 48), (18, -34, 48)  Amygdala |
| (Canal-Rivero et al., 2025) | Psychosis | Gray matter voxel-based morphometry | Precuneus (−6, −70, 38), (−2, −60, 18)  Cuneus (−10, −100, 0), (−14, −104, −6)  Cerebellum (−24, −36, −30), (−24, −42, −24), (−28, −26, −32), (−34, −56, −22)  Amygdala | Cuneus (10, −96, 24)  Amygdala |
| (Liu et al., 2025) | NSSI | Gray matter voxel-based morphometry and resting-state fMRI connectivity | Pallidum (-31.5, -3, 6)  Middle temporal gyrus (-45, -36, 0), (-51, -39, 6)  Angular gyrus (-45, -36, 0) | Pallidum (31.5, -3, 9)  Middle temporal gyrus (48, -27, 3) |
| (Whittle et al., 2009) | NSSI | Gray matter volumetry | ACC |  |
| (Takahashi et al., 2009) | NSSI | Gray matter volumetry |  |  |
| (Takahashi et al., 2010) | NSSI | Gray matter volumetry |  |  |
| (Spoletini et al., 2011) | Psychosis | Gray matter volumetry |  | Amygdala |
| (Giakoumatos et al., 2013) | Psychosis | Gray matter volumetry | Inferior Temporal  Superior Parietal  Supramarginal gyrus  Thalamus | Inferior Temporal  Rostral Middle Frontal  Superior Frontal  Fusiform  Lingual  Nucleus Accumbens |
| (Nanda et al., 2016) | Psychosis | Gray matter volumetry |  |  |
| (Ando et al., 2018) | NSSI | Gray matter volumetry | ACC | ACC |
| (Yin et al., 2023) | Psychosis | Gray matter volumetry | Amygdala  Basal nucleus, accessory basal, cortico-amygdaloid transition, paralaminar nucleus  Hippocampus | Basal nucleus, paralaminar nucleus |
| (Wang et al., 2022) | NSSI | Gray matter volumetry and cortical thickness |  |  |
| (Besteher et al., 2016) | Psychosis | Gray matter cortical thickness |  | Middle temporal (42, 21, - 35)  Temporopolar (42, 21, - 35) |
| (Girgis et al., 2021) | Psychosis | Gray matter cortical thickness | Middle temporal gyrus | Middle temporal gyrus |
| (Huber et al., 2025) | NSSI | Gray matter cortical thickness and volumetry |  | Right rostral ACC |
| (Rüsch et al., 2008) | Psychosis | Gray matter voxel-based morphometry and white matter volumetry | Inferior frontal gyrus (-31, 25, -7) | Inferior frontal gyrus (41, 25, -9) |
| (Duerden et al., 2014) | NSSI | Gray matter cortical thickness and white matter FA & MD | Primary somatosensory cortex (-56, -18, 53) | Superior parietal lobule (11 -51 74)  Primary somatosensory cortex (11, -45, 75) |
| (S.-J. Lee et al., 2016) | Psychosis | White matter FA | Cerebral peduncle  Posterior limb of the internal capsule  Retrolenticular part of the internal capsule  Anterior, superior, and posterior corona radiata  External capsule  Superior longitudinal fasiculus  Posterior thalamic radiation | Posterior thalamic radiation |
| (Long et al., 2018) | Psychosis | White matter FA & MD | Corpus callosum  Anterior, superior, and posterior corona radiata  Posterior thalamic radiation  Superior longitudinal fasciculus | Corpus callosum  Posterior corona radiata |
| (Patel et al., 2023) | NSSI | Gray matter cortical surface area | Inferior parietal cortex (-34.9 -87.8 13.1)  Precentral (-53.0, -2.7, 7.8) |  |
| (Plener et al., 2012) | NSSI | fMRI task activation | Cerebellum (-17, -51, -29), (-31, -76, -50)  Superior frontal cortex (-20, 28, 59)  Superior parietal cortex (-28, -62, 51)  IFG (-31, 32, -16)  Hippocampus (-24, -18, -21)  Amygdala (-24, -18, -21)  ACC (-2, 7, 26)  Middle frontal gyrus (-28, 36, -4)  Cuneus (-13, -87, 21)  Occipital cortex (-24, -47, 30) | Cerebellum (12, -72, -25), (34, -65, -33), (5, -54, -50), (12, -72, -25), (5, -54, -50), (23, -58, -50)  Inferior parietal cortex (70, -33, -4)  IFG (48, 0, 34)  Middle temporal cortex (37, -69, 17)  Hippocampus (16, 0, -25)  Amygdala (16, 0, -25)  ACC (-2, 7, 26)  Middle occipital gyrus (41, -72, 26)  Inferior frontal cortex (59, 32, 13) |
| (Minzenberg et al., 2014) | Psychosis | fMRI task activation | Superior frontal gyrus (-22, 40, 38), (-16, 54, 40), (-26, 50, 40), (-20, 42, 38), (-20, 52,38), (-8, 54, 40)  Medial frontal gyrus (-4, 40, 28), (-12, 44, 14), (-2, 55, 12), (-12, 48, 12), (-8, -14, 62), (-8, 0, 50), (-10, 10, 52)  Middle frontal gyrus (-26, -12, 58), (-30, -8, 50), (-36, 8, 40), (-16, -10, 66)  Inferior frontal gyrus (-42, 8, 10) | Superior frontal gyrus (24, -6, 58), (24, 42, 52)  Middle frontal gyrus (34, 54, 4), (36, 56, 4), (34, 26, 40)  Medial frontal gyrus (6, 50 ,10), (15, 42, 15)  Inferior frontal gyrus (48, 24, 8)  Dorsal anterior cingulate gyrus (10, 0, 44), (15, 4, 42), (16, 8, 40), (15, 4, 42), (6, 46, 12), (4, 32, 30)  Precentral gyrus (38, -8, 38), (48, -5, 32) |
| (Bonenberger et al., 2015) | NSSI | fMRI task activation |  |  |
| (K.-H. Lee et al., 2015) | Psychosis | fMRI task activation |  | DLPFC (16, 42, 31) |
| (Minzenberg et al., 2015a) | Psychosis | fMRI task activation |  | Inferior frontal gyrus (22, 40, 2), (12, 24, 4), (40, 28, -4)  Precentral gyrus (44, 0, 2)  Middle frontal gyrus (24, 50, 14) |
| (Minzenberg et al., 2015b) | Psychosis | fMRI task activation | SMA  Pre-SMA  Premotor cortex  DLPFC |  |
| (Groschwitz et al., 2016) | NSSI | fMRI task activation | Ventrolateral PFC (-30, -28, -12)  Medial PFC (-8, 42, 6)  Parahippocampus (-28, -40, -16) | Medial PFC (12, 52, 8)  Pre-SMA (54, 4, 30) |
| (Sauder et al., 2016) | NSSI | fMRI task activation | Amygdala (-18, -3, -12) | Amygdala (30, -3, -27) |
| (Quevedo et al., 2016) | NSSI | fMRI task activation | Dorsal PFC (-16, 32, 56)  Parahippocampus, fusiform, hippocampus (-30, -34, -12), (-18, -32, -16)  Middle occipital gyrus, lingual gyrus, cuneus (-20, -88, -18)  Rostral medial PFC (-8, 58, 14) | Precuneus (12, -40, 48), (8, -58, 30)  Superior parietal lobule (30, -60, 60)  Parahippocampus, fusiform, hippocampus (28, 2, -28), (-18, -32, -16)  Amygdala (28, 2, -28)  Middle temporal gyrus (58, -60, 0) |
| (Brown et al., 2017) | NSSI | fMRI task activation | DLPFC (-44, 34, 28)  Dorsomedial PFC (-18, 22, 55) | Premotor cortex (36, 0, 40) |
| (Potvin et al., 2018) | Psychosis | fMRI task activation | Middle occipital gyrus (19, -30, 7) | Cerebellar declive (18, -64, -29)  Lingual gyrus (18, -88, -4)  Medial frontal gyrus (19, 47, 8), (19, 50, 11)  ACC (19, 47, 8), (19, 50, 11)  Culmen (21, -64, -32)  Cuneus (18, -80, 33) |
| (Demers et al., 2019) | NSSI | fMRI task activation | Precentral gyrus (-16, -34, 64) | Supramarginal gyrus (64, -44, 26)  Inferior frontal gyrus (46, 12, 24) |
| (Poon et al., 2019) | NSSI | fMRI task activation |  |  |
| (Mayo et al., 2021) | NSSI | fMRI task activation |  |  |
| (Malejko et al., 2022) | NSSI | fMRI task activation | Dorsal ACC (10, 22, 38)  IFG, pars triangularis (-48, 40, 10), (-48, 26, 10)  IFG, pars opercularis (-38, 4, 28)  Supramarginal gyrus (-64, -34, 30)  SMA (2, -4, 56)  Inferior parietal cortex (-42, -32, 40), (-42, -44, 52)  Precuneus (-12, -46, 52) | Dorsal ACC (10, 22, 38)  IFG, pars triangularis (38, 26, 28), (64, 32, 4)  IFG, pars opercularis (44, 16, 30), (50, 14, 6)  Supramarginal gyrus (58, -42, 26)  SMA (2, -4, 56)  Middle frontal gyrus (38, 26, 28) |
| (Westlund Schreiner et al., 2023) | NSSI | fMRI task activation | Temporal-occipital fusiform cortex and lingual gyrus (-24, -50, -10)  Lateral occipital cortex (-8, -70, 64)  Superior lateral occipital cortex (-18, -82, 50)  Hippocampus (-12, -16, -16)  Amygdala (-12, -16, -16)  Precuneus (-8, -70, 64), (-6, -62, 16)  Intracalcarine cortex (-6, -62, 16)  Cerebellum Vermis Crus II (0, -76, -30)  Cerebellum VI (-2, -78, -18)  Superior and middle frontal gyrus (-26, 10, 58) | Cerebellum Crus I (44, -48, -40)  Parahippocampal and lingual gyrus (28, -42, -8)  Middle frontal and precentral gyrus (44, 6, 46) |
| (Hoptman et al., 2024) | Psychosis | fMRI task activation | MFG (-1, 47, 27)  Superior frontal gyrus (-13, 52, 36) | Rostral ACC (12, 37, 13)  Superior frontal gyrus (28, 59, 15), (26, 47, 32)  DLPFC (30, 39, 32) |
| (Athanassiou et al., 2021) | Psychosis | fMRI task activation & connectivity | Middle frontal gyrus (-32, 30, 46)  Precuneus (-2, -54, 54), (-4, 8, 30)  Hippocampus (-14, -40, 6)  Primary motor cortex (-44, -16, 46)  Precentral area (-52, -2, 26)  Rolandic operculum (-46, -22, 14)  Primary motor cortex (-24, -28, 60)  Intracalcarine cortex (-22, -74, 10) | Primary motor cortex (32, -24, 70)  Cerebellum (10, -40, -18)  Intracalcarine cortex (-2, -84, 2)  Superior frontal gyrus (24, 30 ,42) |
| (Osuch et al., 2014) | NSSI | fMRI task seed-based connectivity | Precuneus, cuneus, occipital gyrus, angular gyrus (-8, -72, 14), (-2, -60, 6), (-22, -68, 8)  Culmen, parahippocampal gyrus, lingual gyrus (-16, -30, -8), (-2, -44, -2), (-2, -52, -14), (-2, -38, -12), (-8, -28, -16), (-14, -32, -20)  Cuneus and precuneus (-8, -82, 44) | Precuneus, cuneus, supramarginal gyrus (12, -64, 40), (12, -78, 42), (2, -76, 34), (36, -56, 34), (44, -62, 40), (56, -48, 36), (18, -68, 14)  Thalamus, globus pallidus (10, -24, 30), (8, -20, 20), (22, -10, 20), (8, -18, 8)  Midbrain, pons, parahippocampal gyrus, inferior frontal gyrus, amygdala, culmen (16, -10, -20), (28, 8, -20),  Middle frontal gyrus (18, 34, -16)  Occipital, cuneus, and precuneus (16, -92, 34), (18, -98, 22), (4, -94, 20), (2, -90, 32) |
| (Minzenberg et al., 2016) | Psychosis | fMRI task seed-based connectivity | Dorsal ACC  Middle frontal gyrus (-50, 28, 36), (-54, 28, 36), (-52, 16, 34)  Precentral gyrus (-48, 6, 28), (-52, 6, 26)  Superior frontal gyrus (-24, -2, 44), (-10, 18, 62)  Middle temporal gyrus (-50, -72, 28), (-50, -72, 26)  Superior parietal lobule (-48, -60, 50), (-48, -60, 30), (-56, -56, 28) | Dorsal ACC  Superior frontal gyrus (22, 8, 46), (22, -4, 68)  Middle frontal gyrus (36, 2, 42), (30, 6, 48), (50, 12, 42), (38, 2, 42), (32, 6, 46)  Inferior parietal lobule ( 42, -52, 28), (52, -52, 66), (38, -62, 44), (58, -54, 54)  Middle temporal gyrus (48, -70, 16), (40, -72, 14), (42, -54, -4), (46, -64, -4)  Middle occipital gyrus (42, -82, 18), (28, -94, 12), (36, -92, 4)  Globus pallidus (6, 2, -4)  Inferior frontal gyrus (30, 28, -4)  Postcentral gyrus (22, -36, 80), (32, -46, 72), (30, -44, 72)  Inferior temporal gyrus (52, -74, -2)  Precentral gyrus (26, -18, 68) |
| (L. Lin et al., 2024) | NSSI | fMRI task seed-based connectivity | Precuneus (-12, -4, 44) | SMA (6, -26, 56) |
| (Santana-Gonzalez et al., 2025) | NSSI | fMRI task seed-based connectivity | Amygdala  Ventral striatum  Precuneus (-25, -77, 47)  Inferior parietal lobule, precentral gyrus, postcentral gyrus (-63, -26, 45) | Amygdala  Ventral striatum  Fusiform gyrus, Parahippocampal gyrus (14, -50, -2) |
| (H. Zhang et al., 2013) | Psychosis | fMRI task effective connectivity | Medial PFC (-3, 56, 22) |  |
| (Perini et al., 2019) | NSSI | fMRI task MVPA classification | Superior/middle occipital gyrus (-31, -80, 26), (-37, -81, 1)  Middle temporal gyrus (-56, -14, -19)  Dorsolateral PFC (-50, 40, 14), (-38, 20, 34)  Dorsomedial PFC (-4, 59, 20)  Subgenual ACC (-4, 15, -14)  Precuneus (-25, -70, 30)  Postcentral gyrus (-37, -35, 61)  Cerebellum lobule IX (-14, -41, -47) | Paracentral lobule (8, -38, 62)  Dorsomedial PFC (24, 52, 19), (12, 58, 30)  Dorsolateral PFC (42, 26, 26)  Ventrolateral PFC (30, 49, 11)  Middle temporal gyrus (40, -78, 8)  Parietal operculum (62, -3, 15)  Ventral cuneus (2, -86, 9)  Cerebellum pyramis (8, -67, -35) |
| (Westlund Schreiner et al., 2017) | NSSI | fMRI task and resting state seed-based connectivity | ACC (2, 10, 34), (2, 10, 32)  SMA (2, 10, 34), (2, 10, 32)  Angular gyrus (-44, -56, 34), (-46, -58, 34)  Occipital cortex (-44, -56, 34), (-46, -58, 34), (-32, -42, 38)  Middle temporal gyrus (-58, -26, -8)  Amygdala  Superior parietal lobule (-34, -42, 38)  Frontal pole (0, 56, -4)  Medial frontal cortex (0, 56, -4)  Paracingulate (0, 56, -4) | ACC (2, 10, 34), (2, 10, 32)  SMA (2, 10, 34), (2, 10, 32)  Frontal pole (24, 66, 14), (0, 56, -4)  Inferior temporal gyrus (50, -8, -36)  Middle temporal gyrus (50, -8, -36)  Temporal pole (50, -8, -36)  Angular gyrus (40, -68, 28)  Occipital cortex (40, -68, 28), (32, -66, 52), (32, -60, 62)  Amygdala  Lingual gyrus, occipital pole, occipital fusiform, temporal fusiform (26, -66,-18)  Superior parietal lobule (32, -60, 62), (32, -60, 62)  Medial frontal cortex (0, 56, -4)  Paracingulate (0, 56, -4) |
| (Santamarina-Perez et al., 2019) | NSSI | fMRI resting state seed-based connectivity | Medial PFC  Amygdala | ACC (6, 30, -2)  Subcallosal cortex (6, 30, -2)  Paracingulate (6, 30, -2)  Planum temporale (56, -26, 14)  Precentral gyrus (14, -16, 56)  Postcentral gyrus (14, -16, 56)  Medial PFC  Amygdala |
| (Cullen et al., 2020) | NSSI | fMRI resting state seed-based connectivity | Amygdala  Precuneus  Nucleus accumbens  Medial frontal gyrus | Amygdala  Inferior frontal cortex  Medial frontal gyrus  SMA  Middle frontal gyrus  Middle frontal cortex  Nucleus accumbens |
| (Otto et al., 2023) | NSSI | fMRI resting state seed-based connectivity | Central opercular cortex | Middle frontal gyrus  Superior frontal gyrus  Frontal pole  Posterior parietal cortex  Lateral prefrontal cortex  Angular gyrus  Planum temporale |
| (Choi et al., 2024) | NSSI | fMRI resting state seed-based connectivity | Secondary visual area (-24, -74, -10)  Fusiform gyrus (-24, -74, -10)  Premotor cortex (-14, 0, 48)  SMA (-14, 0, 48) | Dorsolateral PFC (36, 64, 24) |
| (Huang et al., 2021) | NSSI | fMRI resting state ALFF and seed-based connectivity | Pallidum (-12, 3, -3) | Fusiform gyrus (27, -36, -15)  Paracingulate gyrus (12, -24, 39)  Postcentral gyrus (63, -33, 3)  Inferior parietal lobule (45, -39, 57) |
| (Liao et al., 2024) | NSSI | fMRI resting state ALFF and seed-based connectivity | Inferior occipital gyrus (-39, -78, -9)  Middle occipital gyrus (-24, -87, -1)  Lingual gyrus (-9, -78, 0)  Fusiform gyrus (-42, -51, 15)  ACC (0, 36, 30)  Medial superior frontal gyrus (0, 33, 32)  Superior parietal gyrus (-21, -63, -51)  Precuneus (-14, -69, 57)  Rolandic operculum (-48, -1, 3)  Middle temporal gyrus (-60, -31, -13)  Inferior temporal gyrus (-57, -6, -36)  Postcentral gyrus (-54, -18, -45)  Inferior parietal angular gyrus (-53, -18, 41) | Inferior parietal angular gyrus (45, -39, 54)  Superior parietal gyrus (48, -38, 59)  Inferior frontal gyrus (48, 9, 24)  Dorsolateral superior frontal gyrus (27, -9, 63)  Middle frontal gyrus (33, 33, 48)  SMA (6, 15, 51)  Precentral gyrus (42, -9, 57)  Postcentral gyrus (27, -39, 54)  Lingual gyrus (12, -57, 3) |
| (Zhou et al., 2022) | NSSI | fMRI resting state ALFF, fALFF, and ReHo | Middle occipital cortex (-33, -87, 3), (-24, -87, 15), (-36, -69, 39), (-33, -90, 12), (-33, -87, 9)  Calcarine (-9, -96, 0), (-15, -93, 0), (-18, -75, 12)  Supramarginal gyrus (-60, -24, 33), (-60, -27, 27)  Angular gyrus (-42, -63, 42)  Hippocampus (-24, -12, -21), (-24, -36, 6)  Precuneus (-6, -42, 39), (0, -57, 51)  Middle frontal gyrus (-33, 45, 39)  Superior frontal (-15, 54, 39) | Middle temporal cortex (60, -66, 21)  Middle occipital cortex (42, -81, 27)  Superior medial frontal (6, 48, 45)  Superior frontal (24, 45, 42) |
| (X. Lin et al., 2025) | NSSI | fMRI resting state ALFF and resting state functional connectivity | Hippocampus (-21, -24, -3)  Precuneus (-12, -51, 42)  Thalamus (-18, -24, 0)  Dorsal lateral superior frontal gyrus (-9, 45, 48) | Hippocampus (24, -27, 3)  SMA (9, -12, 60)  Middle temporal gyrus (57, -57, 9), (54, -72, 6), (54, -33, 0)  Inferotemporal gyrus (60, -3, -30)  Postcentral gyrus (63, -9, 36)  Superior parietal gyrus (24, -57, 63) |
| (Chen et al., 2023) | NSSI | fMRI resting state functional connectivity | Nucleus accumbens  Middle frontal gyrus (-27, 3, 45)  Superior frontal gyrus (-27, 54, 0)  Inferior cerebellum (-18, -69, -39)  Middle temporal gyrus (-48, -66, 18), (-59, -65, 15)  Thalamus (-21, -27, 9) | Amygdala  Hippocampus  Thalamus  Supramarginal gyrus (51, -27, 39)  Lingual gyrus (15, -39, 12)  Angular gyrus (51, -69, 42)  Superior parietal gyrus (27, -72, 51), (30, -72, 51), (33, -72, 54)  Angular gyrus (30, -72, 51), (54, -63, 24)  Middle temporal gyrus (54, -63, 24), (54, -63, 24)  Rolandic operculum (54, -24, 21)  Superior frontal gyrus (21, -12, 60)  Middle occipital gyrus (30, -84, 21) |
| (Ho et al., 2021) | NSSI | fMRI resting state ICA and seed-to-seed connectivity | Ventral DMN  Anterior DMN | Ventral DMN  Anterior DMN |
| (J. Zhang et al., 2025) | NSSI | fMRI resting state ICA and voxel-wise connectivity | Posterior DMN  Frontoparietal network  Anterior DMN | Posterior DMN  Anterior DMN  Middle frontal gyrus (39, 24, 27)  Angular gyrus (33, -57, 39) |

**Supplemental Table 1b.** Significant ROIs present in literature from adolescents with NSSI and suicidality in psychosis that were not selected for GIMME analysis. Coordinates in MNI space were listed if they were available. FA = fractional anisotropy; MD = mean diffusivity; PFC = prefrontal cortex; IFG = inferior frontal gyrus; MFG = medial frontal gyrus; ACC = anterior cingulate cortex; SMA = supplementary motor area; DLPFC = dorsolateral prefrontal cortex; DMN = default mode network.

| **Cortical Parcel** |  |
| --- | --- |
| R.MCC  17Networks_RH_SalVentAttnA_ParMed_1 | L.MCC  17Networks_LH_SalVentAttnA_FrMed_1 |
| R.PCC  17Networks_RH_DefaultA_PCC_3  17Networks_RH_DefaultA_PCC_4 | L.PCC  17Networks_LH_DefaultA_PCC_2  17Networks_LH_DefaultA_PCC_3  17Networks_LH_DefaultA_PCC_4 |
| R.Caudate  FreeSurfer_Right-Caudate | L.Caudate  FreeSurfer_Left-Caudate |
| R.Putamen  FreeSurfer_Right-Putamen | L.Putamen  FreeSurfer_Left-Putamen |
| R.STG  17Networks_RH_SomMotB_S2_6  17Networks_RH_SomMotB_S2_7 | L.STG  17Networks_LH_SomMotB_Aud_5  17Networks_LH_SomMotB_Aud_6  17Networks_LH_SomMotB_Aud_7 |
| R.OFC  17Networks_RH_Limbic_OFC_1  17Networks_RH_Limbic_OFC_2  17Networks_RH_Limbic_OFC_5 | L.OFC  17Networks_LH_Limbic_OFC_2  17Networks_LH_Limbic_OFC_4 |
| R.Insula  17Networks_RH_SalVentAttnA_Ins_3  17Networks_RH_SalVentAttnA_Ins_4  17Networks_RH_SalVentAttnA_Ins_5  17Networks_RH_SalVentAttnA_Ins_6 | L.Insula  17Networks_LH_SalVentAttnA_Ins_1  17Networks_LH_SalVentAttnA_Ins_3  17Networks_LH_SalVentAttnA_Ins_4  17Networks_LH_SalVentAttnB_PFCv_3 |

**Supplemental Table 2.** Schaefer 400 parcellation atlas regions averaged to create selected ROIs. ROI = region of interest; L = left; R = right; MCC = middle cingulate cortex; PCC = posterior cingulate cortex; STG = superior temporal gyrus; OFC = orbitofrontal cortex.

*2.* *Sensitivity Analysis*

| **Path** | **Path Type** | **Path Stability** | | |
| --- | --- | --- | --- | --- |
|  |  | EP vs. NCC | NSSI- vs. NSSI+ | SA- vs. SA+ |
| L.Caudate → L.Caudate lag | group | 100% | 100% | 100% |
| L.Putamen → L.Caudate | group | 98.24% | 100% | 92.98% |
| R.Putamen → L.Caudate | group | + (5.26%) | NA | + (7.01%) |
| L.Insula → L.Insula lag | group | 100% | 100% | 100% |
| R.STG → L.Insula | group | 100% | 100% | 100% |
| L.Insula → L.MCC | group | + (1.75%) | + (1.75%) | + (1.75%) |
| L.MCC → L.MCC lag | group | 100% | 100% | 100% |
| L.STG → L.MCC | group | 92.98% | 92.98% | 96.49% |
| R.MCC → L.MCC | group | 100% | 100% | 100% |
| R.STG → L.MCC | group | + (5.26%) | NA | + (1.75%) |
| L.OFC → L.OFC lag | group | 100% | 100% | 100% |
| R.OFC → L.OFC | group | + (3.50%) | + (3.50%) | + (3.50%) |
| L.PCC → L.PCC lag | group | 100% | 100% | 100% |
| L.STG → L.PCC | group | 100% | 100% | 100% |
| L.Insula → L.Putamen | group | 100% | 100% | 100% |
| L.Insula → L.Putamen lag | group | 100% | 100% | 100% |
| L.Putamen → L.Putamen lag | group | 100% | 100% | 100% |
| R.Putamen → L.Putamen lag | group | + (10.52%) | + (1.75%) | NA |
| L.MCC → L.STG | group | + (1.75%) | + (1.75%) | + (1.75%) |
| L.STG → L.STG lag | group | 100% | 100% | 100% |
| R.Insula → L.STG | group | + (10.52%) | + (10.52%) | + (10.52%) |
| R.Putamen → L.STG | group | 87.71% | 87.71% | 87.71% |
| L.Caudate → R.Caudate | group | 100% | 100% | 100% |
| R.Caudate → R.Caudate lag | group | 100% | 100% | 100% |
| R.Putamen → R.Caudate | group | 100% | 100% | 100% |
| L.Insula → R.Insula | group | 100% | 100% | 100% |
| .L.Insula → R.Insula lag | group | 100% | 100% | 100% |
| R.Insula → R.Insula lag | group | 100% | 100% | 100% |
| R.STG → R.Insula | group | 100% | 100% | 100% |
| L.Insula → R.MCC | group | 100% | 100% | 100% |
| L.Insula → R.MCC lag | group | 89.47% | 85.96% | 98.24% |
| R.Insula → R.MCC | group | 100% | 100% | 100% |
| R.MCC → R.MCC lag | group | 100% | 100% | 100% |
| L.OFC → R.OFC | group | 96.49% | 50.87% | 50.87% |
| R.OFC → R.OFC lag | group | 100% | 100% | 100% |
| L.Insula → R.PCC | group | + (3.50%) | + (3.50%) | + (3.50%) |
| L.PCC → R.PCC | group | 100% | 100% | 100% |
| L.PCC → R.PCC lag | group | 100% | 100% | 100% |
| R.MCC → R.PCC | group | 96.49% | 96.49% | 96.49% |
| R.PCC → R.PCC lag | group | 100% | 100% | 100% |
| L.Putamen → R.Putamen | group | 100% | 100% | 100% |
| R.Insula → R.Putamen | group | + (14.03%) | + (12.28%) | + (12.28%) |
| R.Insula → R.Putamen lag | group | + (14.03%) | + (12.28%) | + (12.28%) |
| R.Putamen → R.Putamen lag | group | 100% | 100% | 100% |
| L.STG → R.STG | group | 100% | 100% | 100% |
| R.STG → R.STG lag | group | 100% | 100% | 100% |
| .L.Putamen → L.Caudate | subgroup | + (1.75%) | NA | + (7.01%) |
| R.Caudate → L.Caudate lag | subgroup | 98.24% | + (3.50%) | + (5.26%) |
| R.PCC → L.Caudate | subgroup | + (10.52%) | 96.49% | 89.47% |
| R.Putamen → L.Caudate lag | subgroup | + (3.50%) | NA | NA |
| L.MCC → L.Insula | subgroup | NA | + (8.77%) | NA |
| L.PCC → L.Insula lag | subgroup | NA | 73.68% | NA |
| R.STG → L.Insula lag | subgroup | + (5.26%) | NA | NA |
| L.Insula → L.MCC | subgroup | 14 | + (10.52%) | + (3.50%) |
| L.PCC → L.MCC | subgroup | NA | + (1.75%) | NA |
| L.STG → L.MCC | subgroup | NA | + (5.26%) | NA |
| R.STG → L.MCC | subgroup | NA | + (5.26%) | NA |
| R.OFC → L.OFC | subgroup | NA | 26 | NA |
| L.STG → L.PCC lag | subgroup | 100% | NA | NA |
| L.STG → L.Putamen | subgroup | + (10.52%) | + (1.75%) | + (1.75%) |
| R.Putamen → L.Putamen | subgroup | NA | 66.66% | NA |
| R.Putamen → L.Putamen lag | subgroup | 89.47% | + (24.56%) | 100% |
| R.STG → L.Putamen | subgroup | + (1.75%) | + (12.28%) | + (7.01%) |
| L.Putamen → L.STG | subgroup | + (5.26%) | NA | + (1.75%) |
| R.Insula → R.MCC lag | subgroup | + (3.50%) | NA | + (1.75%) |
| L.STG → R.MCC | subgroup | NA | + (10.52%) | NA |
| R.Putamen → R.MCC | subgroup | + (14.03%) | + (21.05%) | NA |
| L.OFC → R.OFC | subgroup | NA | + (29.82%) | + (45.61%) |
| L.Insula → R.Putamen lag | subgroup | NA | + (1.75%) | + (1.75%) |
| R.Insula → R.Putamen | subgroup | 85.96% | 87.71% | 87.71% |
| R.Insula → R.Putamen lag | subgroup | 85.96% | 85.96% | 85.96% |

**Supplemental Table 3.** Path stability measures for all paths in each GIMME model calculated from a leave-one-out inspired sensitivity analysis. Path type identifies whether a path is a group-level (representative of 75% of the entire sample) or subgroup-level (representative of 75% of the subgroup participants) path. Path stability of X% represents a path present in the original model which is recovered in X% of the leave-one-out models. Thus, 100% path stability is an original model connection recovered in all leave-one-out models. + represents a path which is not present in the original model and is generated in X% of the leave-one-out models. L = left; R = right; MCC = middle cingulate cortex; PCC = posterior cingulate cortex; STG = superior temporal gyrus; OFC = orbitofrontal cortex.

*3. Path Coefficients*

| Path | Predictor | β coefficient | stderr | t | p | CI Lower | CI Higher | p (FDR) |
| --- | --- | --- | --- | --- | --- | --- | --- | --- |
| NCC vs. EP | | | | | | | | |
| L.Putamen → R.Putamen | Diagnostic Group | -0.095 | 0.043 | -2.231 | 0.030 | -0.181 | -0.009 | 0.979 |
| L.Putamen → L.Putamen lag | Diagnostic Group | -0.152 | 0.056 | -2.731 | 0.009 | -0.264 | -0.040 | 0.296 |
| L.Caudate → L.Caudate lag | Diagnostic Group | 0.130 | 0.037 | 3.559 | 0.001 | 0.057 | 0.203 | 0.029* |
| L.STG → L.PCC | Diagnostic Group | 0.143 | 0.034 | 4.266 | 8.8E-05 | 0.076 | 0.210 | 0.003* |
| L.Putamen → L.Putamen lag | Age | -0.014 | 0.007 | -2.055 | 0.045 | -0.028 | 0.000 | 0.913 |
| L.STG → L.STG lag | Age | -0.013 | 0.005 | -2.362 | 0.022 | -0.024 | -0.002 | 0.751 |
| L.PCC → L.PCC lag | Age | -0.008 | 0.003 | -2.392 | 0.021 | -0.014 | -0.001 | 0.720 |
| L.Putamen → L.Caudate | Age | -0.012 | 0.005 | -2.454 | 0.018 | -0.022 | -0.002 | 0.636 |
| R.Insula → R.MCC | Sex | -0.069 | 0.031 | -2.209 | 0.032 | -0.132 | -0.006 | 0.975 |
| L.STG → R.STG | FD | 0.608 | 0.298 | 2.040 | 0.047 | 0.009 | 1.207 | 0.988 |
| L.Putamen → L.Caudate | FD | 0.472 | 0.224 | 2.105 | 0.040 | 0.022 | 0.922 | 0.988 |
| L.STG → L.MCC | FD | 0.388 | 0.148 | 2.619 | 0.012 | 0.090 | 0.685 | 0.361 |
| R.STG → L.Insula | FD | 0.854 | 0.309 | 2.767 | 0.008 | 0.234 | 1.474 | 0.253 |
| L.PCC → L.PCC lag | FD | -0.449 | 0.149 | -3.006 | 0.004 | -0.749 | -0.149 | 0.136 |
| R.STG → R.Insula | FD | 0.533 | 0.167 | 3.197 | 0.002 | 0.198 | 0.868 | 0.082 |
| L.Insula → L.Insula lag | FD | -0.852 | 0.230 | -3.706 | 0.001 | -1.314 | -0.390 | 0.018* |
| L.STG → L.PCC | FD | 0.758 | 0.195 | 3.892 | 0.000 | 0.367 | 1.149 | 0.011* |

| **Path** | **Predictor** | **β coefficient** | **stderr** | ***t*** | ***p*** | **CI Lower** | **CI Higher** | ***p* (FDR)** |
| --- | --- | --- | --- | --- | --- | --- | --- | --- |
| ***NSSI- vs. NSSI+*** | | | | | | | | |
| R.Insula → R.MCC | Diagnostic Group | -0.066 | 0.031 | -2.114 | 0.039 | -0.128 | -0.003 | 0.945 |
| L.PCC → L.PCC lag | Diagnostic Group | -0.065 | 0.024 | -2.681 | 0.010 | -0.114 | -0.016 | 0.357 |
| L.Insula → L.Putamen | Age | -0.010 | 0.005 | -2.050 | 0.046 | -0.020 | 0.000 | 0.971 |
| L.STG → L.STG lag | Age | -0.012 | 0.006 | -2.171 | 0.035 | -0.023 | -0.001 | 0.971 |
| L.Putamen → L.Putamen lag | Age | -0.014 | 0.006 | -2.344 | 0.023 | -0.026 | -0.002 | 0.785 |
| R.Putamen → R.Caudate | Age | -0.006 | 0.002 | -2.370 | 0.022 | -0.011 | -0.001 | 0.760 |
| L.PCC → L.PCC lag | Age | -0.007 | 0.003 | -2.393 | 0.020 | -0.013 | -0.001 | 0.738 |
| R.Insula → R.MCC | Sex | -0.065 | 0.028 | -2.306 | 0.025 | -0.121 | -0.008 | 0.911 |
| L.STG → R.STG | FD | 0.651 | 0.298 | 2.180 | 0.034 | 0.051 | 1.250 | 0.982 |
| R.STG → L.Insula | FD | 0.723 | 0.277 | 2.612 | 0.012 | 0.167 | 1.279 | 0.356 |
| L.STG → L.MCC | FD | 0.422 | 0.149 | 2.838 | 0.007 | 0.123 | 0.721 | 0.203 |
| L.Caudate → R.Caudate | FD | 0.687 | 0.230 | 2.993 | 0.004 | 0.226 | 1.148 | 0.137 |
| L.STG → L.PCC | FD | 0.661 | 0.182 | 3.633 | 0.001 | 0.295 | 1.026 | 0.022* |
| L.PCC → L.PCC lag | FD | -0.519 | 0.141 | -3.682 | 0.001 | -0.802 | -0.236 | 0.019* |
| L.Insula → L.Insula lag | FD | -0.915 | 0.212 | -4.311 | 7.6E-05 | -1.341 | -0.488 | 0.003* |
| R.STG → R.Insula | FD | 0.611 | 0.159 | 3.837 | 3.5E-04 | 0.291 | 0.932 | 0.012* |
| L.Insula → L.Putamen lag | NSSI | 0.179 | 0.048 | 3.733 | 4.8E-04 | 0.083 | 0.275 | 0.016* |
| L.Insula → L.Putamen | NSSI | -0.171 | 0.044 | -3.916 | 2.7E-04 | -0.259 | -0.083 | 0.009* |
| L.Putamen → L.Putamen lag | NSSI | -0.214 | 0.053 | -4.075 | 1.6E-04 | -0.320 | -0.109 | 0.006* |
| R.Putamen → R.Putamen lag | NSSI | 0.173 | 0.038 | 4.550 | 3.4E-05 | 0.097 | 0.250 | 0.001* |
| L.Putamen → R.Putamen | NSSI | -0.260 | 0.052 | -4.996 | 7.5E-06 | -0.365 | -0.156 | 2.7E-04* |

| **Path** | **Predictor** | **β coefficient** | **stderr** | ***t*** | ***p*** | **CI Lower** | **CI Higher** | ***p* (FDR)** |
| --- | --- | --- | --- | --- | --- | --- | --- | --- |
| ***SA- vs. SA+*** | | | | | | | | |
| L.PCC → L.PCC lag | Diagnostic Group | -0.063 | 0.024 | -2.587 | 0.013 | -0.112 | -0.014 | 0.454 |
| L.STG → L.STG lag | Age | -0.012 | 0.006 | -2.075 | 0.043 | -0.023 | 0.000 | 0.963 |
| R.Putamen → R.Caudate | Age | -0.005 | 0.002 | -2.243 | 0.029 | -0.010 | -0.001 | 0.963 |
| L.PCC → L.PCC lag | Age | -0.007 | 0.003 | -2.439 | 0.018 | -0.014 | -0.001 | 0.660 |
| R.Insula → R.MCC | Sex | -0.060 | 0.029 | -2.109 | 0.040 | -0.118 | -0.003 | 0.981 |
| L.STG → L.MCC | FD | 0.382 | 0.150 | 2.541 | 0.014 | 0.080 | 0.683 | 0.426 |
| R.STG → L.Insula | FD | 0.827 | 0.309 | 2.678 | 0.010 | 0.207 | 1.447 | 0.310 |
| R.STG → R.Insula | FD | 0.512 | 0.167 | 3.060 | 0.004 | 0.176 | 0.849 | 0.114 |
| L.Caudate → R.Caudate | FD | 0.710 | 0.231 | 3.073 | 0.003 | 0.246 | 1.174 | 0.113 |
| L.PCC → L.PCC lag | FD | -0.512 | 0.142 | -3.608 | 0.001 | -0.796 | -0.227 | 0.024* |
| L.Insula → L.Insula lag | FD | -0.860 | 0.234 | -3.682 | 0.001 | -1.330 | -0.391 | 0.020* |
| L.STG → L.PCC | FD | 0.708 | 0.187 | 3.776 | 4.2E-04 | 0.331 | 1.084 | 0.015* |

**Supplemental Table 4.** Significant path coefficients for all three GIMME models 1) non-clinical control [NCC, n=34] vs. early psychosis [EP, n=23], 2) no lifetime presence of non-suicidal self-injury [NSSI-, n=36] vs. lifetime presence of non-suicidal self-injury [NSSI+, n=21], 3) no lifetime history of suicide attempt(s) [SA-, n=36] vs. lifetime history of suicide attempt(s) [SA+, n=21]. All paths are group-level paths from each respective model. Statistics were calculated from a regression including diagnostic group, NSSI presence, number of SA, age, sex, and mean FD. stderr = standard error; CI = confidence interval; FDR = false discovery rate corrected; * = significant at FDR corrected p < 0.05; FD = framewise displacement; L = left; R = right; MCC = middle cingulate cortex; PCC = posterior cingulate cortex; STG = superior temporal gyrus; OFC = orbitofrontal cortex.

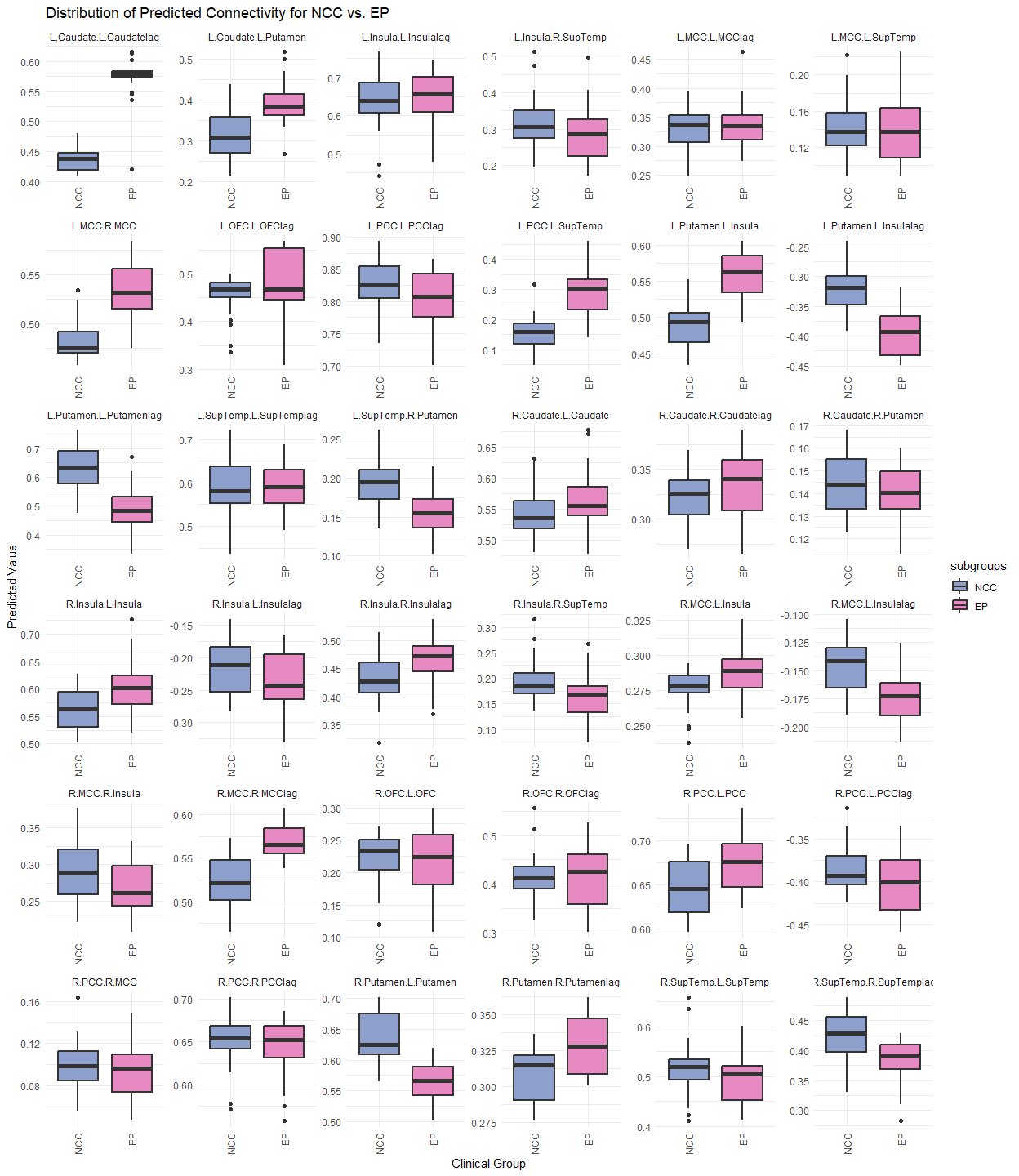
*4. Predicted Effective Connectivity Strength*

**Supplemental Figure 1.** Distribution of predicted effective connectivity strength for non-clinical controls [NCC] (left, n=34) compared to participants with early psychosis [EP] (right, n=23). All paths are group-level paths from the NCC vs. EP model. Predicted effective connectivity strength was calculated from a regression including diagnostic group, NSSI presence, number of SA, age, sex, and mean FD. Periods in this case represent connectivity from the region on the right to the region on the left (←). Red dots represent the median. L = left; R = right; MCC = middle cingulate cortex; PCC = posterior cingulate cortex; SupTemp = superior temporal gyrus; OFC = orbitofrontal cortex.

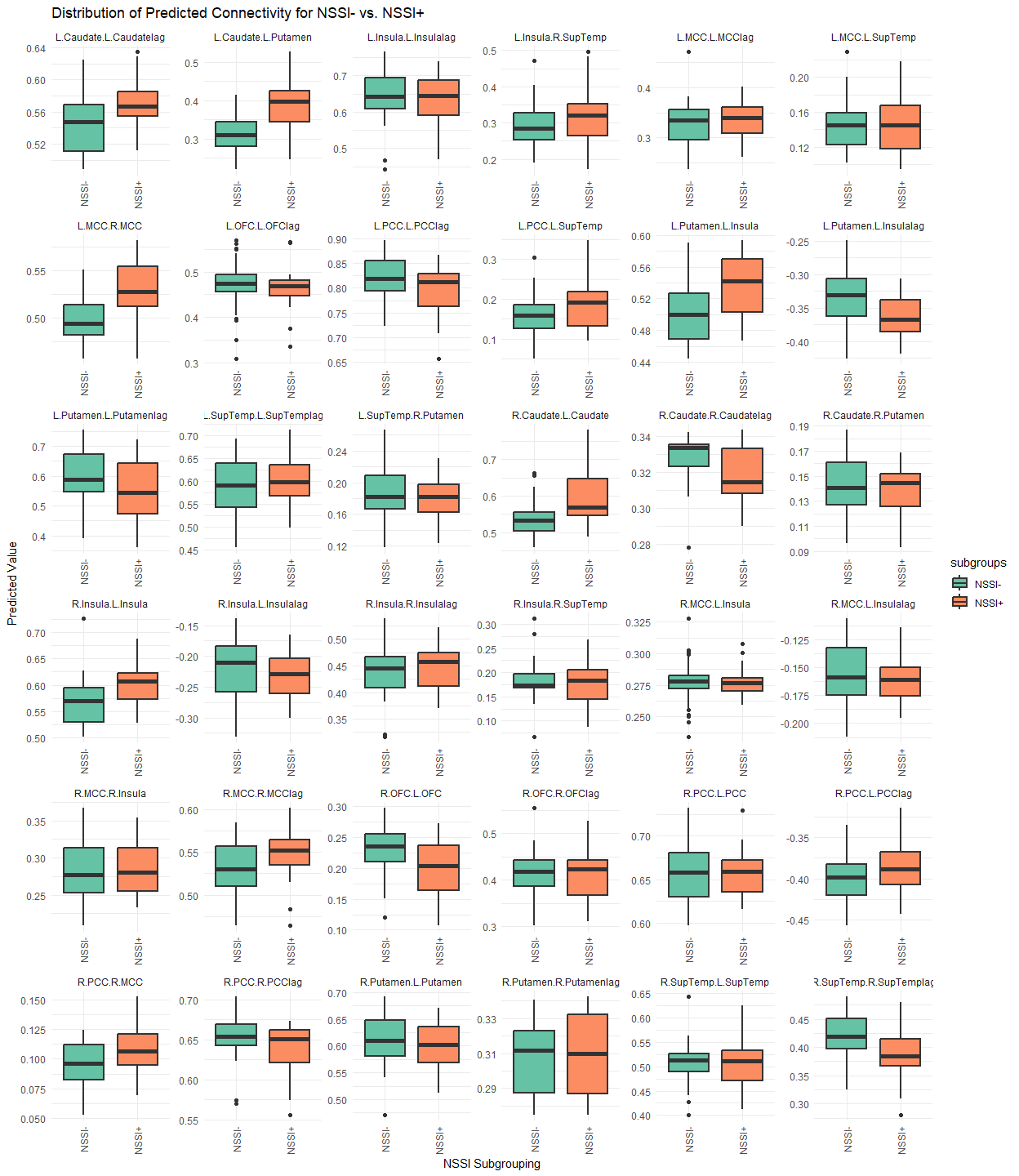

**Supplemental Figure 2.** Distribution of predicted effective connectivity strength for participants without lifetime history of non-suicidal self-injury [NSSI-] (left, n=36) compared to participants with lifetime history of non-suicidal self-injury [NSSI+] (right, n=21). All paths are group-level paths from the NSSI- vs. NSSI+ model. Predicted effective connectivity strength was calculated from a regression including diagnostic group, NSSI presence, number of SA, age, sex, and mean FD. Periods in this case represent connectivity from the region on the right to the region on the left (←). Red dots represent the median. L = left; R = right; MCC = middle cingulate cortex; PCC = posterior cingulate cortex; SupTemp = superior temporal gyrus; OFC = orbitofrontal cortex.

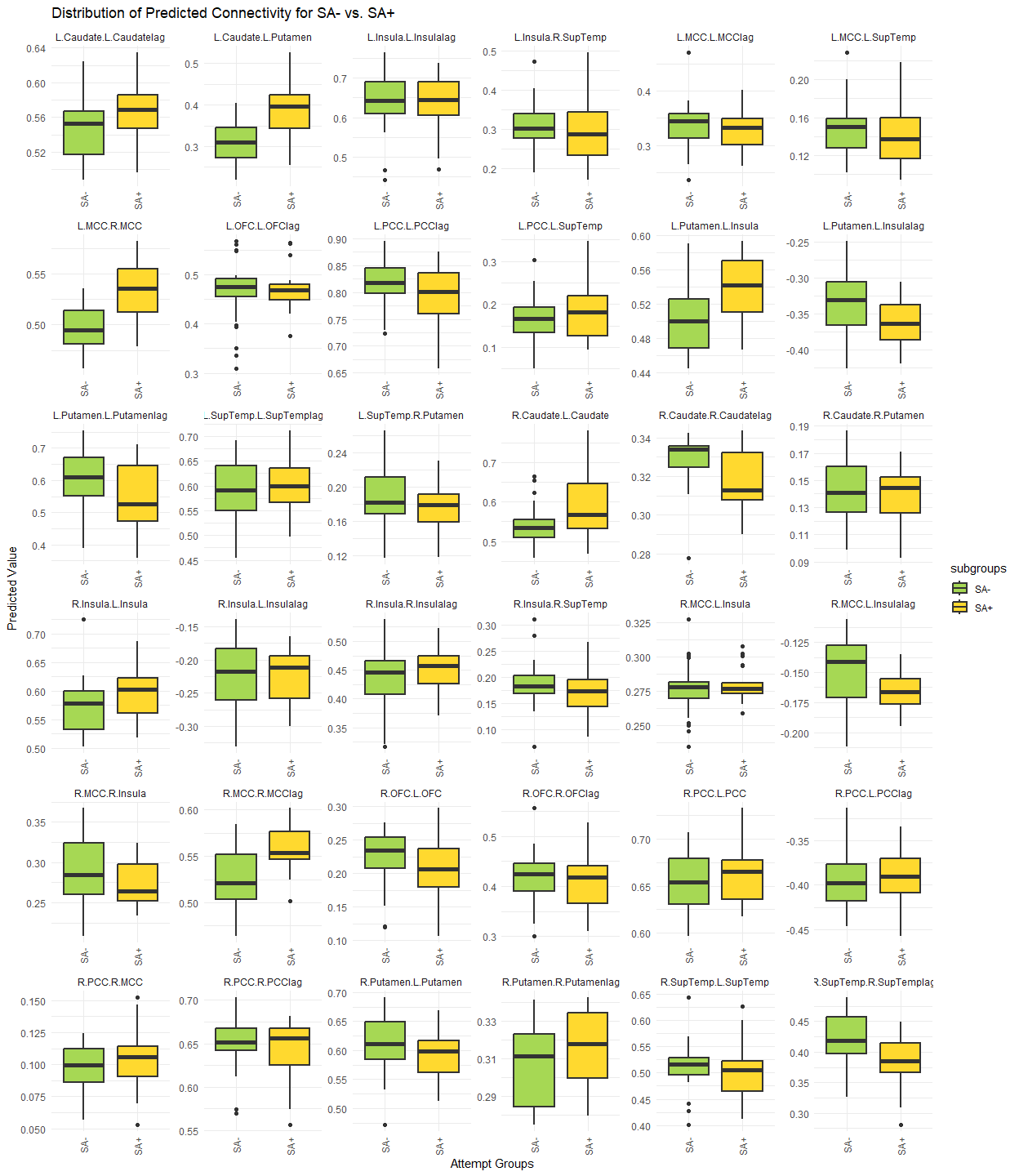

**Supplemental Figure 3.** Distribution of predicted effective connectivity strength for participants without lifetime history of suicide attempt(s) [SA-] (left, n=36) compared to participants with lifetime history of suicide attempt(s) [SA+] (right, n=21). All paths are group-level paths from the SA- vs. SA+ model. Predicted effective connectivity strength was calculated from a regression including diagnostic group, NSSI presence, number of SA, age, sex, and mean FD. Periods in this case represent connectivity from the region on the right to the region on the left (←). Red dots represent the median. L = left; R = right; MCC = middle cingulate cortex; PCC = posterior cingulate cortex; SupTemp = superior temporal gyrus; OFC = orbitofrontal cortex.

**References**

Aguilar, E. J., García-Martí, G., Martí-Bonmatí, L., Lull, J. J., Moratal, D., Escartí, M. J., Robles, M., González, J. C., Guillamón, M. I., & Sanjuán, J. (2008). Left orbitofrontal and superior temporal gyrus structural changes associated to suicidal behavior in patients with schizophrenia. *Progress in Neuro-Psychopharmacology and Biological Psychiatry*, *32*(7), 1673–1676. https://doi.org/10.1016/j.pnpbp.2008.06.016

Ando, A., Reichl, C., Scheu, F., Bykova, A., Parzer, P., Resch, F., Brunner, R., & Kaess, M. (2018). Regional grey matter volume reduction in adolescents engaging in non-suicidal self-injury. *Psychiatry Research: Neuroimaging*, *280*, 48–55. https://doi.org/10.1016/j.pscychresns.2018.08.005

Athanassiou, M., Dumais, A., Iammatteo, V., De Benedictis, L., Dubreucq, J.-L., & Potvin, S. (2021). The processing of angry faces in schizophrenia patients with a history of suicide: An fMRI study examining brain activity and connectivity. *Progress in Neuro-Psychopharmacology and Biological Psychiatry*, *107*, 110253. https://doi.org/10.1016/j.pnpbp.2021.110253

Beauchaine, T. P., Sauder, C. L., Derbidge, C. M., & Uyeji, L. L. (2019). Self-injuring Adolescent Girls Exhibit Insular Cortex Volumetric Abnormalities that are Similar to those Seen in Adults with Borderline Personality Disorder. *Development and Psychopathology*, *31*(4), 1203–1212. https://doi.org/10.1017/S0954579418000822

Besteher, B., Wagner, G., Koch, K., Schachtzabel, C., Reichenbach, J. R., Schlösser, R., Sauer, H., & Schultz, C. C. (2016). Pronounced prefronto-temporal cortical thinning in schizophrenia: Neuroanatomical correlate of suicidal behavior? *Schizophrenia Research*, *176*(2), 151–157. https://doi.org/10.1016/j.schres.2016.08.010

Bonenberger, M., Plener, P. L., Groschwitz, R. C., Grön, G., & Abler, B. (2015). Differential neural processing of unpleasant haptic sensations in somatic and affective partitions of the insula in non-suicidal self-injury (NSSI). *Psychiatry Research: Neuroimaging*, *234*(3), 298–304. https://doi.org/10.1016/j.pscychresns.2015.10.013

Brown, R. C., Plener, P. L., Groen, G., Neff, D., Bonenberger, M., & Abler, B. (2017). Differential Neural Processing of Social Exclusion and Inclusion in Adolescents with Non-Suicidal Self-Injury and Young Adults with Borderline Personality Disorder. *Frontiers in Psychiatry*, *8*. https://doi.org/10.3389/fpsyt.2017.00267

Canal-Rivero, M., Tordesillas-Gutiérrez, D., Ruiz-Veguilla, M., Ortiz-García de la Foz, V., Cuevas-Esteban, J., Marco de Lucas, E., Vázquez-Bourgon, J., Ayesa-Arriola, R., & Crespo-Facorro, B. (2020). Brain grey matter abnormalities in first episode non-affective psychosis patients with suicidal behaviours: The role of neurocognitive functioning. *Progress in Neuro-Psychopharmacology and Biological Psychiatry*, *102*, 109948. https://doi.org/10.1016/j.pnpbp.2020.109948

Canal-Rivero, M., Tordesillas-Gutiérrez, D., Ruiz-Veguilla, M., Ortiz-García de la Foz, V., Marco de Lucas, E., Romero-Garcia, R., Vázquez-Bourgon, J., Ayesa-Arriola, R., & Crespo-Facorro, B. (2025). Suicidal Behaviour Prior to First Episode Psychosis: Wider and More Widespread Grey-Matter Alterations. *Archives of Suicide Research: Official Journal of the International Academy for Suicide Research*, 1–15. https://doi.org/10.1080/13811118.2025.2454581

Chen, X., Chen, H., Liu, J., Tang, H., Zhou, J., Liu, P., Tian, Y., Wang, X., Lu, F., & Zhou, J. (2023). Functional connectivity alterations in reward-related circuits associated with non-suicidal self-injury behaviors in drug-naïve adolescents with depression. *Journal of Psychiatric Research*, *163*, 270–277. https://doi.org/10.1016/j.jpsychires.2023.05.068

Choi, E. J., Vandewouw, M. M., Taylor, M. J., Stevenson, R. A., Arnold, P. D., Brian, J., Crosbie, J., Kelley, E., Liu, X., Jones, J., Lai, M.-C., Schachar, R. J., Lerch, J. P., & Anagnostou, E. (2024). Dorsal Striatal Functional Connectivity and Repetitive Behavior Dimensions in Children and Youths With Neurodevelopmental Disorders. *Biological Psychiatry: Cognitive Neuroscience and Neuroimaging*, *9*(4), 387–397. https://doi.org/10.1016/j.bpsc.2023.10.014

Cullen, K. R., Westlund Schreiner, M., Klimes-Dougan, B., Eberly, L. E., LaRiviere, L. L., Lim, K. O., Camchong, J., & Mueller, B. A. (2020). Neural correlates of clinical improvement in response to N-acetylcysteine in adolescents with non-suicidal self-injury. *Progress in Neuro-Psychopharmacology and Biological Psychiatry*, *99*, 109778. https://doi.org/10.1016/j.pnpbp.2019.109778

Demers, L. A., Westlund Schreiner, M., Hunt, R. H., Mueller, B. A., Klimes-Dougan, B., Thomas, K. M., & Cullen, K. R. (2019). Alexithymia is associated with neural reactivity to masked emotional faces in adolescents who self-harm. *Journal of Affective Disorders*, *249*, 253–261. https://doi.org/10.1016/j.jad.2019.02.038

Duerden, E. G., Card, D., Roberts, S. W., Mak-Fan, K. M., Chakravarty, M. M., Lerch, J. P., & Taylor, M. J. (2014). Self-injurious behaviours are associated with alterations in the somatosensory system in children with autism spectrum disorder. *Brain Structure and Function*, *219*(4), 1251–1261. https://doi.org/10.1007/s00429-013-0562-2

Giakoumatos, C. I., Tandon, N., Shah, J., Mathew, I. T., Brady, R. O., Clementz, B. A., Pearlson, G. D., Thaker, G. K., Tamminga, C. A., Sweeney, J. A., & Keshavan, M. S. (2013). Are Structural Brain Abnormalities Associated With Suicidal Behavior In Patients With Psychotic Disorders? *Journal of Psychiatric Research*, *47*(10), 1389–1395. https://doi.org/10.1016/j.jpsychires.2013.06.011

Girgis, R. R. (2020). The neurobiology of suicide in psychosis: A systematic review. *Journal of Psychopharmacology*, *34*(8), 811–819. https://doi.org/10.1177/0269881120936919

Girgis, R. R., Basavaraju, R., France, J., Wall, M. M., Brucato, G., Lieberman, J. A., & Provenzano, F. A. (2021). An Exploratory Magnetic Resonance Imaging Study of Suicidal Ideation in Individuals at Clinical High-Risk for Psychosis. *Psychiatry Research. Neuroimaging*, *312*, 111287. https://doi.org/10.1016/j.pscychresns.2021.111287

Groschwitz, R. C., Plener, P. L., Groen, G., Bonenberger, M., & Abler, B. (2016). Differential neural processing of social exclusion in adolescents with non-suicidal self-injury: An fMRI study. *Psychiatry Research: Neuroimaging*, *255*, 43–49. https://doi.org/10.1016/j.pscychresns.2016.08.001

Ho, T. C., Walker, J. C., Teresi, G. I., Kulla, A., Kirshenbaum, J. S., Gifuni, A. J., Singh, M. K., & Gotlib, I. H. (2021). Default mode and salience network alterations in suicidal and non-suicidal self-injurious thoughts and behaviors in adolescents with depression. *Translational Psychiatry*, *11*(1), 38. https://doi.org/10.1038/s41398-020-01103-x

Hoptman, M. J., Evans, K. T., Parincu, Z., Sparpana, A. M., Sullivan, E. F., Ahmed, A. O., & Iosifescu, D. V. (2024). Emotion-related impulsivity and suicidal ideation and behavior in schizophrenia spectrum disorder: A pilot fMRI study. *Frontiers in Psychiatry*, *15*. https://doi.org/10.3389/fpsyt.2024.1408083

Huang, Q., Xiao, M., Ai, M., Chen, J., Wang, W., Hu, L., Cao, J., Wang, M., & Kuang, L. (2021). Disruption of Neural Activity and Functional Connectivity in Adolescents With Major Depressive Disorder Who Engage in Non-suicidal Self-Injury: A Resting-State fMRI Study. *Frontiers in Psychiatry*, *12*. https://doi.org/10.3389/fpsyt.2021.571532

Huber, R. S., Subramaniam, P., Heinrich, L., Boxer, D. J., Shi, X., Schreiner, M. W., Renshaw, P. F., Yurgelun-Todd, D. A., & Kondo, D. G. (2025). Cingulate cortex cortical thickness associated with non-suicidal self-injury and suicide risk in youth with mood disorders. *Journal of Affective Disorders*, *381*, 518–524. https://doi.org/10.1016/j.jad.2025.04.047

Lee, K.-H., Pluck, G., Lekka, N., Horton, A., Wilkinson, I. D., & Woodruff, P. W. R. (2015). Self-harm in schizophrenia is associated with dorsolateral prefrontal and posterior cingulate activity. *Progress in Neuro-Psychopharmacology and Biological Psychiatry*, *61*, 18–23. https://doi.org/10.1016/j.pnpbp.2015.03.005

Lee, S.-J., Kim, B., Oh, D., Kim, M.-K., Kim, K.-H., Bang, S. Y., Choi, T. K., & Lee, S.-H. (2016). White matter alterations associated with suicide in patients with schizophrenia or schizophreniform disorder. *Psychiatry Research: Neuroimaging*, *248*, 23–29. https://doi.org/10.1016/j.pscychresns.2016.01.011

Liao, K., Yu, R., Chen, Y., Chen, X., Wu, X., Huang, X., & Liu, N. (2024). Alterations of regional brain activity and corresponding brain circuits in drug-naïve adolescents with nonsuicidal self-injury. *Scientific Reports*, *14*(1), 24997. https://doi.org/10.1038/s41598-024-75714-5

Lin, L., Liu, Y., Qiu, S., Yang, Y., Yang, Y., Tian, M., Wang, S., Zhang, J., Bai, X., & Xu, Z. (2024). Orbital frontal cortex functional connectivity during gain anticipation linking the rumination and non-suicidal self-injury in late adolescence. *Journal of Affective Disorders*, *350*, 673–680. https://doi.org/10.1016/j.jad.2024.01.117

Lin, X., Sun, Y., Hu, Y., Chen, Y., Gu, Y., Chen, K., Zou, Z., Li, Z., Wu, X., Li, T., Lv, X., Yan, F., Dou, S., Guo, Z., Chang, S., & Li, Y. (2025). Impaired neural activity and functional connectivity in the hippocampus of adolescents with non-suicidal self-injury addiction. *BMC Psychiatry*, *25*, 895. https://doi.org/10.1186/s12888-025-07331-z

Liu, X., Zhang, Y., Chen, J., Xie, M., Pan, L., Hommel, B., Yang, Y., Zhu, X., Wang, K., & Zhang, W. (2025). Altered brain structure and function correlate with non-suicidal self-injury in children and adolescents with transdiagnostic psychiatric disorders. *Journal of Psychiatric Research*, *184*, 17–26. https://doi.org/10.1016/j.jpsychires.2025.02.051

Long, Y., Ouyang, X., Liu, Z., Chen, X., Hu, X., Lee, E., Chen, E. Y. H., Pu, W., Shan, B., & Rohrbaugh, R. M. (2018). Associations Among Suicidal Ideation, White Matter Integrity and Cognitive Deficit in First-Episode Schizophrenia. *Frontiers in Psychiatry*, *9*. https://doi.org/10.3389/fpsyt.2018.00391

Malejko, K., Hafner, S., Brown, R. C., Plener, P. L., Grön, G., Graf, H., & Abler, B. (2022). Neural Signatures of Error Processing in Depressed Adolescents with Comorbid Non-Suicidal Self-Injury (NSSI). *Biomedicines*, *10*(12), 3188. https://doi.org/10.3390/biomedicines10123188

Mayo, L. M., Perini, I., Gustafsson, P. A., Hamilton, J. P., Kämpe, R., Heilig, M., & Zetterqvist, M. (2021). Psychophysiological and Neural Support for Enhanced Emotional Reactivity in Female Adolescents With Nonsuicidal Self-injury. *Biological Psychiatry: Cognitive Neuroscience and Neuroimaging*, *6*(7), 682–691. https://doi.org/10.1016/j.bpsc.2020.11.004

Minzenberg, M. J., Lesh, T. A., Niendam, T. A., Cheng, Y., & Carter, C. S. (2016). Conflict-Related Anterior Cingulate Functional Connectivity Is Associated With Past Suicidal Ideation and Behavior in Recent-Onset Psychotic Major Mood Disorders. *The Journal of Neuropsychiatry and Clinical Neurosciences*, *28*(4), 299–305. https://doi.org/10.1176/appi.neuropsych.15120422

Minzenberg, M. J., Lesh, T. A., Niendam, T. A., Yoon, J. H., Cheng, Y., Rhoades, R. N., & Carter, C. S. (2015a). Control-related frontal-striatal function is associated with past suicidal ideation and behavior in patients with recent-onset psychotic major mood disorders. *Journal of Affective Disorders*, *188*, 202–209. https://doi.org/10.1016/j.jad.2015.08.049

Minzenberg, M. J., Lesh, T. A., Niendam, T. A., Yoon, J. H., Rhoades, R. N., & Carter, C. S. (2014). Frontal cortex control dysfunction related to long-term suicide risk in recent-onset schizophrenia. *Schizophrenia Research*, *157*(1), 19–25. https://doi.org/10.1016/j.schres.2014.05.039

Minzenberg, M. J., Lesh, T., Niendam, T., Yoon, J. H., Cheng, Y., Rhoades, R. N., & Carter, C. S. (2015b). Frontal Motor Cortex Activity During Reactive Control Is Associated With Past Suicidal Behavior in Recent-Onset Schizophrenia. *Crisis*, *36*(5), 363–370. https://doi.org/10.1027/0227-5910/a000335

Nanda, P., Tandon, N., Mathew, I. T., Padmanabhan, J. L., Clementz, B. A., Pearlson, G. D., Sweeney, J. A., Tamminga, C. A., & Keshavan, M. S. (2016). Impulsivity across the psychosis spectrum: Correlates of cortical volume, suicidal history, and social and global function. *Schizophrenia Research*, *170*(1), 80–86. https://doi.org/10.1016/j.schres.2015.11.030

Osuch, E., Ford, K., Wrath, A., Bartha, R., & Neufeld, R. (2014). Functional MRI of pain application in youth who engaged in repetitive non-suicidal self-injury vs. Psychiatric controls. *Psychiatry Research: Neuroimaging*, *223*(2), 104–112. https://doi.org/10.1016/j.pscychresns.2014.05.003

Otto, A., Jarvers, I., Kandsperger, S., Reichl, C., Ando, A., Koenig, J., Kaess, M., & Brunner, R. (2023). Stress-induced alterations in resting-state functional connectivity among adolescents with non-suicidal self-injury. *Journal of Affective Disorders*, *339*, 162–171. https://doi.org/10.1016/j.jad.2023.07.032

Patel, K. K., Sheridan, M. A., Bonar, A. S., Giletta, M., Hastings, P. D., Nock, M. K., Rudolph, K. D., Slavich, G. M., Prinstein, M. J., & Miller, A. B. (2023). A preliminary investigation into cortical structural alterations in adolescents with nonsuicidal self-injury. *Psychiatry Research: Neuroimaging*, *336*, 111725. https://doi.org/10.1016/j.pscychresns.2023.111725

Perini, I., Gustafsson, P. A., Hamilton, J. P., Kämpe, R., Mayo, L. M., Heilig, M., & Zetterqvist, M. (2019). Brain-based Classification of Negative Social Bias in Adolescents With Nonsuicidal Self-injury: Findings From Simulated Online Social Interaction. *EClinicalMedicine*, *13*, 81–90. https://doi.org/10.1016/j.eclinm.2019.06.016

Plener, P. L., Bubalo, N., Fladung, A. K., Ludolph, A. G., & Lulé, D. (2012). Prone to excitement: Adolescent females with non-suicidal self-injury (NSSI) show altered cortical pattern to emotional and NSS-related material. *Psychiatry Research: Neuroimaging*, *203*(2), 146–152. https://doi.org/10.1016/j.pscychresns.2011.12.012

Poon, J. A., Thompson, J. C., Forbes, E. E., & Chaplin, T. M. (2019). Adolescents’ Reward-related Neural Activation: Links to Thoughts of Nonsuicidal Self-Injury. *Suicide and Life-Threatening Behavior*, *49*(1), 76–89. https://doi.org/10.1111/sltb.12418

Potvin, S., Tikàsz, A., Richard-Devantoy, S., Lungu, O., & Dumais, A. (2018). History of Suicide Attempt Is Associated with Reduced Medial Prefrontal Cortex Activity during Emotional Decision-Making among Men with Schizophrenia: An Exploratory fMRI Study. *Schizophrenia Research and Treatment*, *2018*, 9898654. https://doi.org/10.1155/2018/9898654

Quevedo, K., Martin, J., Scott, H., Smyda, G., & Pfeifer, J. H. (2016). The neurobiology of self-knowledge in depressed and self-injurious youth. *Psychiatry Research: Neuroimaging*, *254*, 145–155. https://doi.org/10.1016/j.pscychresns.2016.06.015

Rüsch, N., Spoletini, I., Wilke, M., Martinotti, G., Bria, P., Trequattrini, A., Bonaviri, G., Caltagirone, C., & Spalletta, G. (2008). Inferior frontal white matter volume and suicidality in schizophrenia. *Psychiatry Research: Neuroimaging*, *164*(3), 206–214. https://doi.org/10.1016/j.pscychresns.2007.12.011

Santamarina-Perez, P., Romero, S., Mendez, I., Leslie, S. M., Packer, M. M., Sugranyes, G., Picado, M., Font, E., Moreno, E., Martinez, E., Morer, A., Romero, M., & Singh, M. K. (2019). Fronto-Limbic Connectivity as a Predictor of Improvement in Nonsuicidal Self-Injury in Adolescents Following Psychotherapy. *Journal of Child and Adolescent Psychopharmacology*, *29*(6), 456–465. https://doi.org/10.1089/cap.2018.0152

Santana-Gonzalez, C., Ranatunga, J., Nguyen, G., Greiskalns, B., Das, N., Lattimer, E., Maurice, M., Yi, G., Zietlow, A.-L., Eckstein, M., Zilverstand, A., & Quevedo, K. (2025). Emotion regulation in self-injurious youth: A tale of two circuits. *Psychiatry Research: Neuroimaging*, *347*, 111944. https://doi.org/10.1016/j.pscychresns.2024.111944

Sauder, C. L., Derbidge, C. M., & Beauchaine, T. P. (2016). Neural responses to monetary incentives among self-injuring adolescent girls. *Development and Psychopathology*, *28*(1), 277–291. https://doi.org/10.1017/S0954579415000449

Spoletini, I., Piras, F., Fagioli, S., Rubino, I. A., Martinotti, G., Siracusano, A., Caltagirone, C., & Spalletta, G. (2011). Suicidal attempts and increased right amygdala volume in schizophrenia. *Schizophrenia Research*, *125*(1), 30–40. https://doi.org/10.1016/j.schres.2010.08.023

Takahashi, T., Chanen, A. M., Wood, S. J., Yücel, M., Kawasaki, Y., McGorry, P. D., Suzuki, M., Velakoulis, D., & Pantelis, C. (2010). Superior temporal gyrus volume in teenagers with first-presentation borderline personality disorder. *Psychiatry Research: Neuroimaging*, *182*(1), 73–76. https://doi.org/10.1016/j.pscychresns.2009.10.014

Takahashi, T., Chanen, A. M., Wood, S. J., Yücel, M., Tanino, R., Suzuki, M., Velakoulis, D., Pantelis, C., & McGorry, P. D. (2009). Insular cortex volume and impulsivity in teenagers with first-presentation borderline personality disorder. *Progress in Neuro-Psychopharmacology and Biological Psychiatry, Bed Nucleus of the Stria Terminalis: Anatomy, Physiology, Functions*, *33*(8), 1395–1400. https://doi.org/10.1016/j.pnpbp.2009.07.017

Wang, K., He, Q., Zhu, X., Hu, Y., Yao, Y., Hommel, B., Beste, C., Liu, J., Yang, Y., & Zhang, W. (2022). Smaller putamen volumes are associated with greater problems in external emotional regulation in depressed adolescents with nonsuicidal self-injury. *Journal of Psychiatric Research*, *155*, 338–346. https://doi.org/10.1016/j.jpsychires.2022.09.014

Westlund Schreiner, M., Klimes-Dougan, B., Mueller, B. A., Eberly, L. E., Reigstad, K. M., Carstedt, P. A., Thomas, K. M., Hunt, R. H., Lim, K. O., & Cullen, K. R. (2017). Multi-modal neuroimaging of adolescents with non-suicidal self-injury: Amygdala functional connectivity. *Journal of Affective Disorders*, *221*, 47–55. https://doi.org/10.1016/j.jad.2017.06.004

Westlund Schreiner, M., Roberts, H., Dillahunt, A. K., Farstead, B., Feldman, D., Thomas, L., Jacobs, R. H., Bessette, K. L., Welsh, R. C., Watkins, E. R., Langenecker, S. A., & Crowell, S. E. (2023). Negative association between non-suicidal self-injury in adolescents and default mode network activation during the distraction blocks of a rumination task. *Suicide and Life-Threatening Behavior*, *53*(3), 510–521. https://doi.org/10.1111/sltb.12960

Whittle, S., Chanen, A. M., Fornito, A., McGorry, P. D., Pantelis, C., & Yücel, M. (2009). Anterior cingulate volume in adolescents with first-presentation borderline personality disorder. *Psychiatry Research: Neuroimaging*, *172*(2), 155–160. https://doi.org/10.1016/j.pscychresns.2008.12.004

Yin, Y., Tong, J., Huang, J., Wang, L., Tian, B., Chen, S., Tan, S., Wang, Z., Yu, T., Li, Y., Tong, Y., Fan, F., Kochunov, P., Hong, L. E., & Tan, Y. (2023). History of suicide attempt associated with amygdala and hippocampus changes among individuals with schizophrenia. *European Archives of Psychiatry and Clinical Neuroscience*, *273*(4), 921–930. https://doi.org/10.1007/s00406-023-01554-5

Zhang, H., Wei, X., Tao, H., Mwansisya, T. E., Pu, W., He, Z., Hu, A., Xu, L., Liu, Z., Shan, B., & Xue, Z. (2013). Opposite Effective Connectivity in the Posterior Cingulate and Medial Prefrontal Cortex between First-Episode Schizophrenic Patients with Suicide Risk and Healthy Controls. *PLOS ONE*, *8*(5), e63477. https://doi.org/10.1371/journal.pone.0063477

Zhang, J., Wu, D., Wang, H., Yu, Y., Zhao, Y., Zheng, H., Wang, S., Fan, S., Pang, X., Wang, K., & Tian, Y. (2025). Large-scale functional network connectivity alterations in adolescents with major depression and non-suicidal self-injury. *Behavioural Brain Research*, *482*, 115443. https://doi.org/10.1016/j.bbr.2025.115443

Zhou, Y., Yu, R., Ai, M., Cao, J., Li, X., Hong, S., Huang, Q., Dai, L., Wang, L., Zhao, L., Zhang, Q., Shi, L., & Kuang, L. (2022). A Resting State Functional Magnetic Resonance Imaging Study of Unmedicated Adolescents With Non-suicidal Self-Injury Behaviors: Evidence From the Amplitude of Low-Frequency Fluctuation and Regional Homogeneity Indicator. *Frontiers in Psychiatry*, *13*. https://doi.org/10.3389/fpsyt.2022.925672
